# Multiple paths to cold tolerance: the role of environmental cues, morphological traits and the circadian clock gene *vrille*

**DOI:** 10.1101/2020.04.20.050351

**Authors:** Noora Poikela, Venera Tyukmaeva, Anneli Hoikkala, Maaria Kankare

**Affiliations:** Department of Biological and Environmental Science, University of Jyväskylä, Finland; Centre d’Ecologie Fonctionelle et Evolutive, CNRS, Montpellier, France

**Author notes:** Corresponding author: Noora Poikela, Department of Biological and Environmental Science, P.O. Box 35, FI-40014 University of Jyväskylä, Finland.

**Keywords:** CT_min_, CCRT, body colour, body weight, latitude, bioclimatic variables, RNA interference (RNAi), *Drosophila montana*, *Drosophila flavomontana*

## Abstract

**Background:** Tracing the association between insect cold tolerance and latitudinally and locally varying environmental conditions, as well as key morphological traits and molecular mechanisms, is essential for understanding the processes involved in adaptation. We explored these issues in two closely-related species, *Drosophila montana* and *Drosophila flavomontana*, originating from diverse climatic locations across several latitudes on the coastal and mountainous regions of North America. We also investigated the association between sequence variation in one of the key circadian clock genes, *vrille*, and cold tolerance in both species. Finally, we studied the impact of *vrille* on fly cold tolerance and cold acclimation ability by silencing it with RNA interference in *D. montana*.

**Results:** We performed a principal component analysis (PCA) on variables representing bioclimatic conditions on the study sites and used latitude as a proxy of photoperiod. PC1 separated the mountainous continental sites from the coastal ones based on temperature variability and precipitation, while PC2 arranged the sites based on summer and annual mean temperatures. Cold tolerance tests showed *D. montana* to be more cold-tolerant than *D. flavomontana* and chill coma resistance (CT_min_) of this species showed an association with PC2. Chill coma recovery time (CCRT) of both species improved towards northern latitudes, and in *D. flavomontana* this trait was also associated with PC1. *D. flavomontana* flies were darkest in the coast and in the northern mountainous populations, but coloration showed no linkage with cold tolerance. Body size decreased towards cold environments in both species, but only within *D. montana* populations largest flies showed fastest recovery from cold. Finally, both the sequence analysis and RNAi study on *vrille* suggested this gene to play an essential role in *D. montana* cold resistance and acclimation, but not in recovery time.

**Conclusions:** Our study demonstrates the complexity of insect cold tolerance and emphasizes the need to trace its association with multiple environmental variables and morphological traits to identify potential agents of natural selection. It also shows that a circadian clock gene *vrille* is essential both for short- and long-term cold acclimation, potentially elucidating the connection between circadian clock system and cold tolerance.

## Background

Species geographical distribution is largely defined by their ability to tolerate stressful conditions and to respond to daily and seasonal temperature changes (e.g. [1-6]). Accordingly, ectothermic species, especially the ones living at high latitudes, may use a variety of physiological and behavioural strategies to increase their cold tolerance [7, 8]. To understand species’ cold adaptation in these regions, it is essential to identify latitudinal and bioclimatic selection pressures driving these adaptations [2, 9–11] and to trace associations between cold tolerance and adaptationally important morphological traits. Also, it is also important to discover molecular mechanisms underlying cold tolerance, but a functional link between candidate genes and cold tolerance has relatively rarely been established (but see e.g. [12, 13]).

Insect cold tolerance is often measured with two ecologically relevant methods. Insects’ chill coma resistance can be assessed by measuring their critical thermal minimum (or chill coma temperature, CT_min_), where they lose neuromuscular coordination and fall into chill coma during a gradual decrease in temperature [14]. Chill coma recovery time (CCRT), on the other hand, is based on time required for an individual to recover from the coma after removal of the chill coma inducing temperature [15, 16]. Even though the ability to resist chill coma and recover from it are thought to be somewhat of a continuum, they are affected by different physiological and molecular mechanisms (e.g. [17, 18]). The onset of chill coma is associated with depolarisation of muscle resting membrane potential due to low temperature (e.g. [19, 20]), while the recovery process involves energetically costly restoration of ion gradients and upregulation of genes involved in repairing cold injuries [21-25]. Both the high resistance to chill coma and the fast recovery from it have several implications on insect fitness in cold environments [15, 18, 26]. For example, fast recovery from chill coma after winter, night or otherwise sudden temperature drops ensures finding high-quality mates on feeding and breeding sites, as well as fast escape from predators. High chill coma resistance covers largely the same advantages, but it may also save insects from falling into energetically costly chill coma. Thus, these traits are likely to be under strong selection.

Latitudinal clines in cold tolerance traits, detected in a variety of species, including *Drosophila* flies [16, 27, 28], *Myrmica* ants [29], *Porcellio laevis* woodlouse [30], highlight adaptive genetic variation in these traits. However, it is challenging to distinguish whether such variation has evolved in response to changes in photoperiod (day length), temperature or their combination [7]. One possible approach is to use GIS (geographic information system) -based environmental data to trace correlations between evolutionary changes in the studied traits and the environments the populations or species experience [31]. Studies on clinal variation in insect cold tolerance can also be complicated by the fact that many species spend the coldest time of the year in reproductive diapause, which induces various kinds of changes in their physiology, metabolism and behaviour in addition to increasing their cold tolerance [32]. Thus, it is important to investigate genetic variation in cold tolerance in temperature and light conditions, where insects’ reproductive status is controlled.

In addition to cold tolerance, morphological traits, like body colour (the degree of melanism) and body size, often show latitudinal variation. Thus, to understand insect adaptation to cold environments, it would be important to investigate cold tolerance traits together with other adaptationally important traits and consider correlations between them. In insects, an increase in melanism towards higher latitudes has been detected both between and within species [33–35]. According to thermal melanin hypothesis, this can be explained by an increased ability of dark individuals to absorb solar radiation and warm up fast in cold environments with low solar radiation (reviewed in [36]). However, body colour may also be affected by other selection pressures, including protection against UV-radiation [37, 38], desiccation [39, 40] and pathogens [41]. Body size is one of the most important quantitative traits associated with insect metabolism, fecundity, mating success and stress tolerance, and thus it is likely under several selection pressures in nature (e.g. [42, 43]). Body size of ectothermic species has been either shown to increase (Bergmann’s rule) or decrease (converse Bergmann’s rule or U-shaped cline) towards higher latitudes and cooler climates (e.g. [42, 44, 45]). Converse Bergmann’s rule applies especially for insects with a long generation time, for which the short growing season on high latitudes limits the time available for development, growth and foraging [45-47]. Overall, parallel clines in cold tolerance and morphological traits give only indirect evidence on the functional linkages between these traits. One way to obtain more direct evidence on their association would be to measure cold tolerance and morphological traits for the same individuals. For example, it has been shown that melanistic wood tiger moths (*Parasemia plantaginis*) warm up more quickly than the less melanistic ones [48] and *Drosophila montana* males’ overwinter survival increases along with an increase in body size in nature [49].

At high latitudes with clear seasonal and diurnal temperature variation, insects can enhance their cold tolerance through long- and short-term cold acclimation [50, 51]. In nature, insects can anticipate the forthcoming cold season from a decreasing temperature and/or day length and adapt to these changes through a gradual increase in cold tolerance [52, 53]. Insects can also be experimentally cold-acclimated by maintaining them in relatively low temperature or short day conditions for a few days to weeks prior to cold shock [54, 55]. Short-term cold acclimation of minutes to hours (rapid cold hardening, RCH) has been suggested to share mechanisms with longer-term acclimation [56] and allow organisms to cope with sudden cold snaps and to optimize their performance during diurnal cooling cycles [24, 53]. Here the circadian clock system, which monitors daily and seasonal light and temperature cycles and entrains behavioural and physiological rhythms to match with them [57], could play a vital role.

In the central circadian clock, described most thoroughly in *D. melanogaster*, daily rhythms are driven via one or several transcriptional feedback loops involving changes in the expression of core circadian clock genes (reviewed in [58-60]). Intriguingly, in *Drosophila montana* flies cold tolerance traits (CT_min_ and CCRT) are under photoperiodic regulation [55, 61], and hence could be associated with circadian clock system. Also, the expression levels of circadian clock genes have been found to change during long-term cold acclimation e.g. in several *Drosophila* species [62-65] and *Gryllus pennsylvanicus* cricket [66]. In plants, the linkage between circadian clock and cold acclimation has been established [67, 68], but in insects direct evidence on this link and its extent is still missing.

Northern *Drosophila virilis* group species possess a very high cold tolerance compared to most other species of the genus [1, 2]. Our study species, *D. montana* and *D. flavomontana*, belong to this group, and they live in diverse climatic conditions in the low-altitude western coast and in the high-altitude Rocky Mountains of North America across several latitudes (Fig. 1). *D. montana* lives generally at higher latitudes and altitudes than *D. flavomontana*, but in some sites the species occur sympatrically [69-71]. Moreover, the body colour of *D. montana* is almost black, while that of *D. flavomontana* varies from light to dark brown [69]. These features make *D. montana* and *D. flavomontana* an ideal species pair to investigate genetic adaptation to cold environments, complementing latitudinal studies performed on less cold-tolerant southern species. They also enable us to study connections between fly cold tolerance and morphological traits and to trace selection pressures underlying cold adaptation. Another interesting point is that the circadian clock system of *D. virilis* group species differs from that of *D. melanogaster*, and shows features that have helped them to adapt to high latitudes [72-76]. Here *vrille*, which is one of the core genes in the central circadian clock system and a key regulator of circadian behavioural rhythms in *D. melanogaster* [58-60], is of special interest, as it has been found to be highly upregulated during cold acclimation in *D. montana* [64, 65, 77].

**Figure 1.**
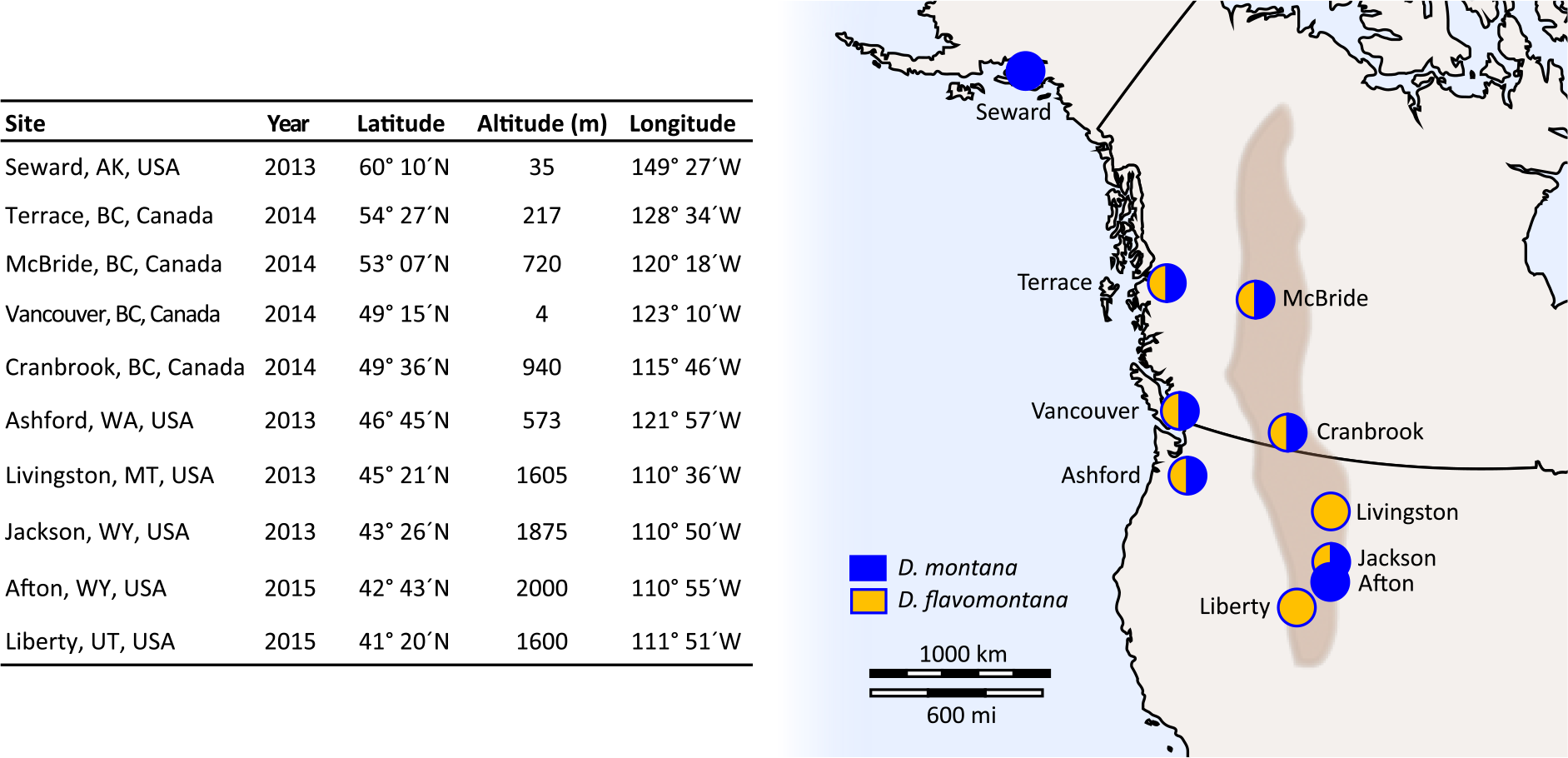
Fly collection sites. Table shows fly collecting sites and years, and the coordinates for each site. Map contains information on whether we have samples from one or both species in each site on the western coast and in the Rocky Mountains (brown area on the map) in North America (detailed information given in Table S1). The map template obtained from https://d-maps.com/carte.php?num_car=5082&lang=en

In this study, we addressed two main objectives. 1) We traced genetic variation in cold tolerance traits (CT_min_ and CCRT) across distribution ranges of *D. montana* and *D. flavomontana*, and identified environmental selection pressures and morphological (body colour and size) correlations driving these adaptations. 2) We compared *vrille* sequence variation to cold tolerance traits among clinal populations of *D. montana* and *D. flavomontana,* and tested the role of this gene in cold tolerance and cold acclimation ability in *D. montana* using RNA interference (RNAi). In the first part of the study, we quantified environmental variation across fly collecting sites by performing a principal component analysis (PCA) on several bioclimatic variables, and investigated whether latitude, as a proxy of photoperiod, and latitudinally or locally varying climatic conditions have shaped fly cold tolerance (CT_min_ and CCRT). Here we predicted that genetic variation in cold tolerance traits shows association with latitudinally varying photoperiods, which serves as the most reliable cue for seasonal temperature changes at given localities [7]. We also quantified genetic variation in fly body colour and body size across species’ distribution range, identified the likely selection pressures affecting these traits and measured direct correlations between the morphological and cold tolerance traits from the same individuals. Body colour could be associated with cold adaptation if it shows latitudinal cline, becoming darker towards northern latitudes, and if it correlates directly with cold tolerance measures (thermal melanin hypothesis, [36]). If large body size and better cold tolerance are associated with each other (e.g [49]), the size could be expected to increase towards cold environments (Bergmann’s rule, [47, 78]) and the traits should be directly correlated. We studied these adaptations in non-diapausing individuals in a common environment to eliminate plastic responses. In the second part of the study, we evaluated the adaptive role of *vrille* in fly cold tolerance by comparing its nucleotide and amino acid variation with variation in two cold tolerance traits (CT_min_ and CCRT) among *D. montana* and *D. flavomontana* cline populations. We also investigated the role of this gene in regulating females’ cold tolerance traits and cold acclimation ability by silencing it with RNAi in *D. montana*. We expected *vrille* to be involved at least in females’ ability to become cold-acclimated, as it shows expression changes during cold acclimation in *D. montana* [64, 77].

## Results

### Variation in the climatic conditions at fly collecting sites

We investigated macroclimatic variability among the sites, where *D. montana* and *D. flavomontana* strains originated from, by performing a principal component analysis (PCA) on 19 bioclimatic variables, growing season length (days) and altitude (Table S2, S3). PCA revealed four principal components (PCs) with eigenvalues > 1 (Table S4). The first two PCs explained more than 83% of the total variation (Fig. 2; Table S4) and were included in further analyses.

**Figure 2.**
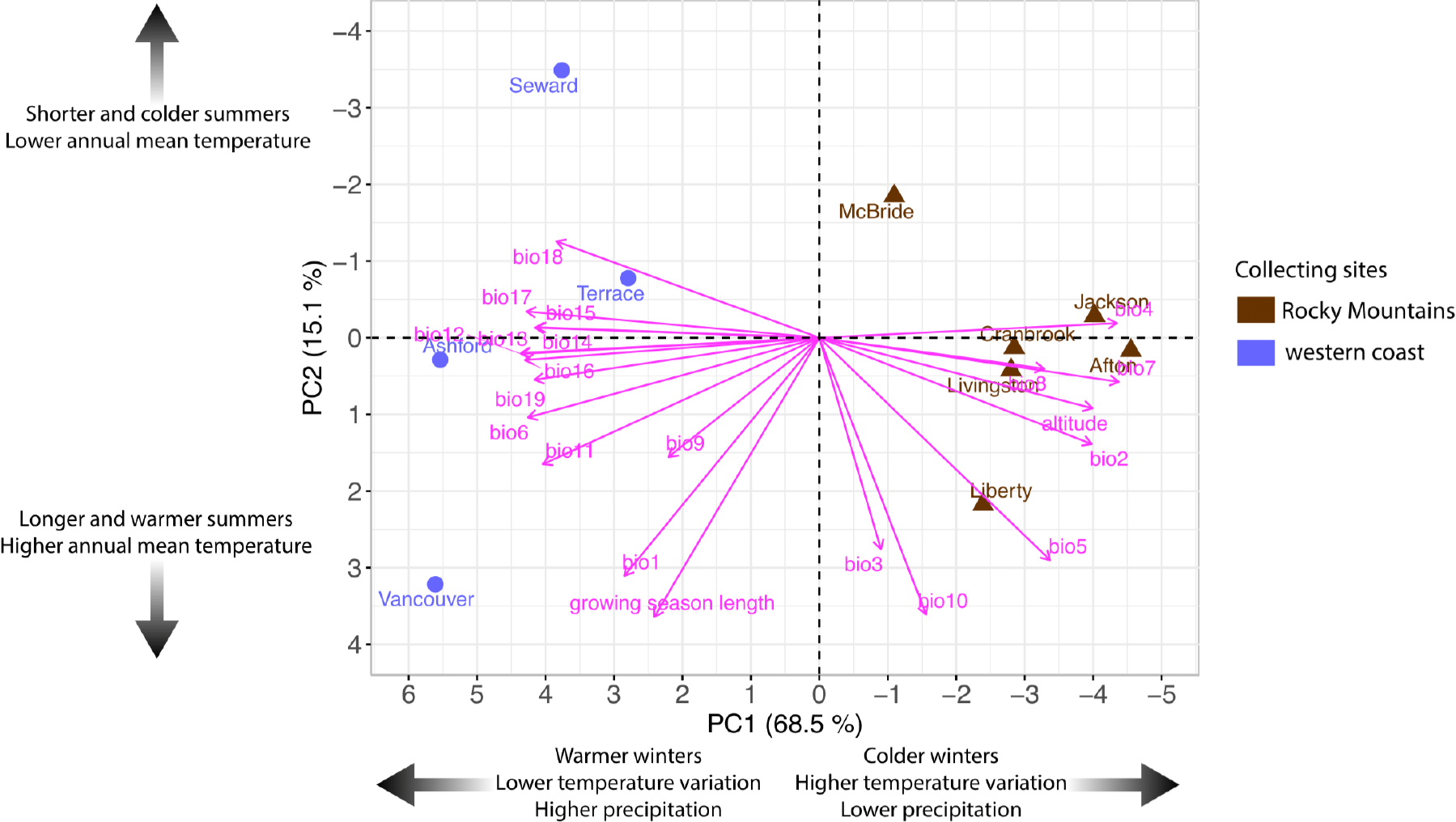
Climatic conditions in the fly collecting sites. Principal component analysis (PCA) was performed on 21 variables describing environmental conditions in fly collecting sites (Table S2, S3). Big black arrows show the change in given conditions.

PC1 clearly separated the Rocky Mountains sites from the ones on the western coast. Variables with the highest contribution on this separation included altitude and the ones describing daily and seasonal temperature variation (bio2, bio4, bio7), the minimum temperature of the coldest month (bio6), and the mean temperature of the coldest quarter (bio11) and precipitation (Fig. 2; Table S5). Together they showed the high-altitude Rocky Mountains sites to have higher temperature variation and colder winters than the ones on the western coast sites. On the other hand, the western coast sites had higher precipitation throughout the year than the Rocky Mountains sites.

PC2 arranged the sites on the basis of the growing season length, annual mean temperature (bio1), the mean temperature of the warmest quarter (bio10), the maximum temperature of the warmest month (bio5) and isothermality (bio3, i.e. how large day-to-night temperatures oscillate relative to the summer-to-winter oscillations; Fig. 2; Table S5). They showed that while some fly collecting sites differ in latitude (photoperiod), they resemble each other in growing season length and summer temperatures due to their variability in altitude and closeness to sea.

### The effects of latitude, climatic conditions and morphological traits on fly cold tolerance

Studying the effects of photoperiod, temperature-related factors (PC1, PC2), body colour and size (measured as weight) on fly cold tolerance enabled us to identify selection pressures affecting this trait. In these experiments, we used three isofemale strains for majority of the populations, but for four populations we succeeded to establish only one or two strains due to the rarity of the species at the collection sites (Table S1). All traits were measured for non-diapausing *D. flavomontana* and *D. montana* females, reared in constant light at 19°C, to minimize plastic responses. Contrary to other traits, weight was analysed only for the flies used in CCRT tests (see Methods). The simplest model, which enabled us to distinguish between latitudinally varying photoperiods and temperatures, included latitude and PC2 as explanatory factors (see Fig. 2; Table S6). The more complicated models included macroclimatic conditions varying between the western coast and the Rocky Mountains (PC1), different interaction terms and fly body colour and size (Table S6).

The best-fit models explaining the cold tolerance (CT_min_ = chill coma temperature, CCRT = chill coma recovery time), body colour and body size of *D. flavomontana* and *D. montana* females are presented in Fig. 3A-D (model comparisons are given in Table S6). The models show that the selection pressures driving the evolution of these traits vary between the species. Most of the pairwise correlations between fly cold tolerance traits, colour and weight were non-significant, and were not included in the best-fit models; Fig. 3, 4; Table S6).

**Figure 3.**
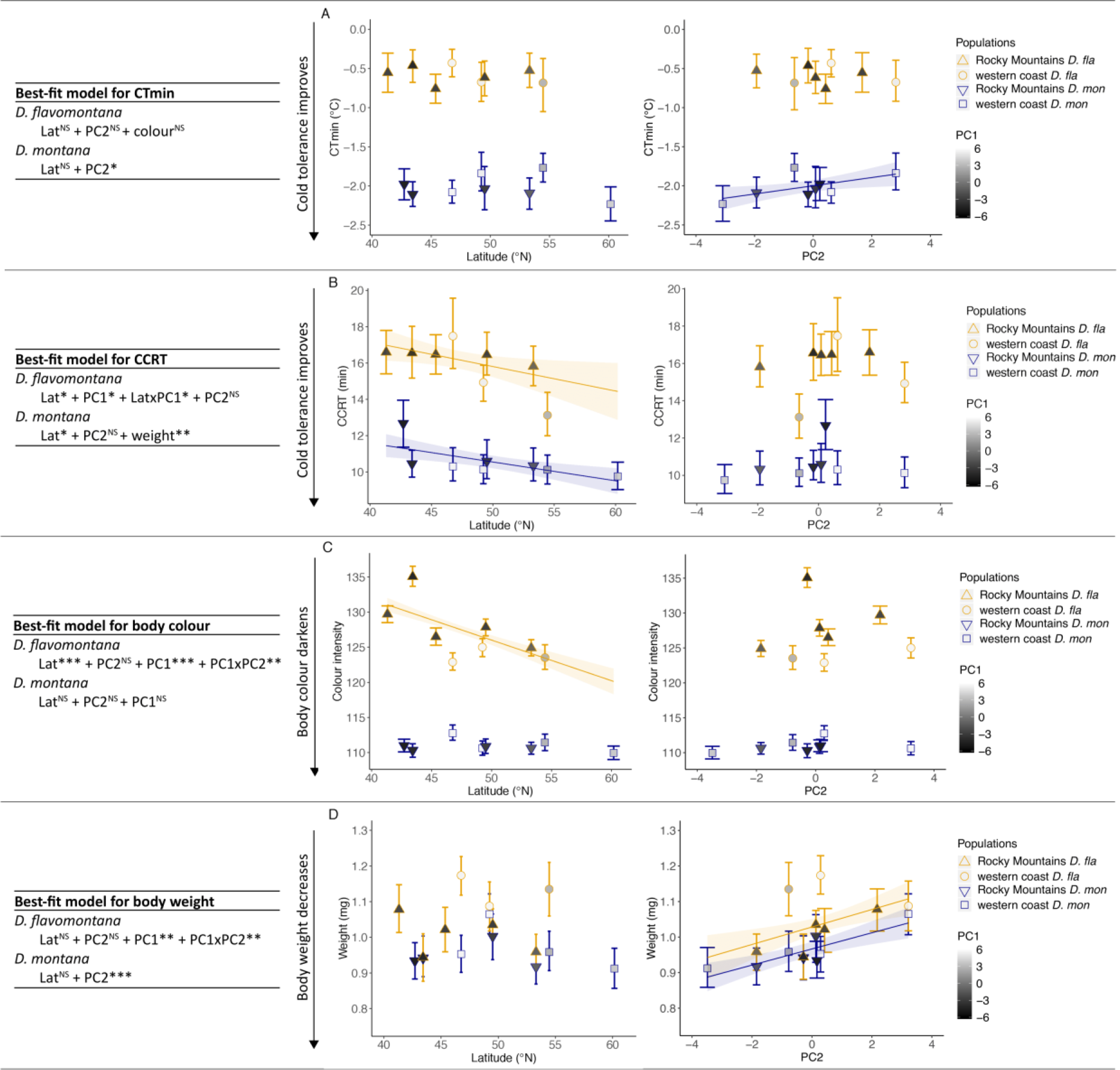
Cold tolerance, morphological traits (body colour and body size) and environmental conditions. The best-fit models are presented in tables. Relationship between latitude (as a proxy of photoperiod) and PC2 (as a proxy of latitudinally varying temperature) and (A) chill coma temperature (CT_min_), (B) chill coma recovery time (CCRT), (C) fly body colour (measured as colour intensity) and (D) fly body size (measured as weight) in *D. flavomontana* (*D. fla*) and *D. montana* (*D. mon*) populations. The effects of local climatic conditions (PC1; see Fig. 2) on the western coast and in the Rocky Mountains are illustrated in grey scale (lighter colours represent the western coast populations and darker ones the Rocky Mountains populations). Error bars represent bootstrapped 95% confidence intervals (Mean ± CI). Significant regression lines for latitude or PC2 with standard errors are shown. Significance levels were obtained from GLMMs or LMM: ^NS^ non-significant, * P < 0.05, ** P < 0.01 and *** P < 0.001.

**Figure 4.**
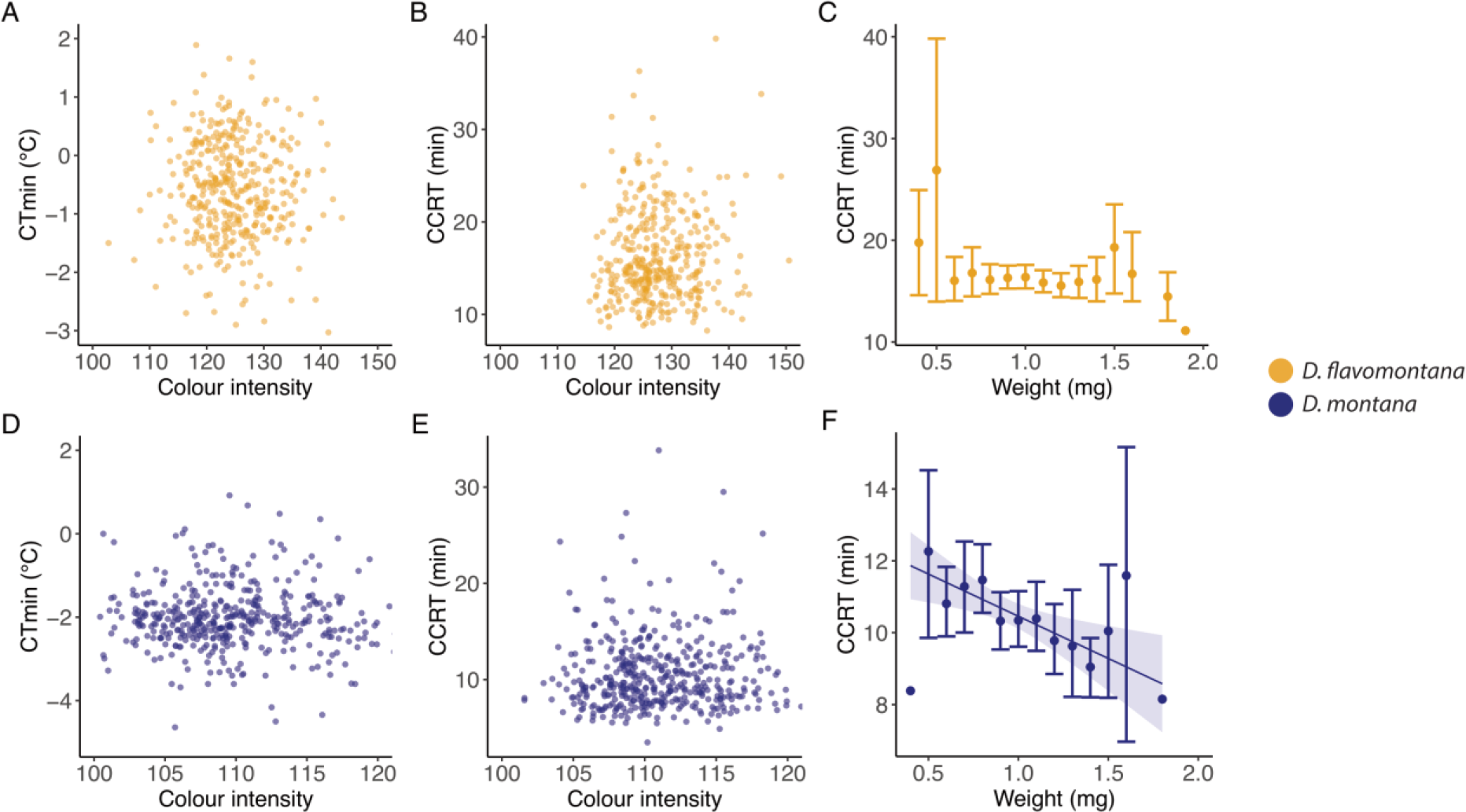
Correlations between fly cold tolerance and morphological traits (body colour or body size). Correlation between (A) *D. flavomontana* CT_min_ and body colour, (B) *D. flavomontana* CCRT and body colour, (C) *D. flavomontana* CCRT and body size (measured as weight), (D) *D. montana* CT_min_ and body colour, (E) *D. montana* CCRT and body colour, (F) *D. montana* CCRT and body size (measured as weight). Error bars represent bootstrapped 95% confidence intervals. Significant regression lines are shown with standard errors (best-fit models are presented in Fig. 3 and Table S6).

CT_min_ of *D. flavomontana* showed only low variation and was not significantly explained by any of the variables (Fig. 3A; Table S7). However, in *D. montana* this trait showed significant association with PC2 (Fig. 3A; Table S7), which suggests that the chill coma resistance of *D. montana* flies is highest in northern populations with a short growing season and cold summer and low annual mean temperatures (Fig. 2). CCRT tests showed *D. flavomontana* flies’ cold tolerance to be significantly associated with latitude and to improve towards North (Fig. 2, 3B; Table S7). Moreover, this trait was affected by PC1, especially on latitudes around 50-55 °N, suggesting that fly cold tolerance is higher in the humid, low-altitude western coast populations than in the high-altitude Rocky Mountains populations with colder temperatures and higher temperature variation (Fig. 2, 3B; Table S7). CCRT of *D. montana* was significantly associated with latitude, improving towards North (Fig. 3B; Table S7). Moreover, in this species large flies recovered faster from chill coma than the small ones (Fig. 3B, 4G; Table S7).

*D. flavomontana* flies’ body colour was significantly affected by latitude, PC1 and an interaction between PC2 and PC1 (Fig. 3C; Table S7). In the Rocky Mountains, the flies became darker (their colour intensity decreased) towards North, while in the western coast populations flies were equally dark and darker than the ones from the Rocky Mountains populations on similar latitudes. *D. montana* body colour showed only minor population-level variation and no significant association with latitude, PC2 or PC1. The body size of *D. flavomontana* was significantly associated with PC1 and an interaction between PC1 and PC2 (Fig. 3D; Table S7), increasing towards warmer winters and summers and longer growing season. The body size of *D. montana* was significantly associated with PC2, being highest in sites with high summer temperatures and a long growing season (Fig. 3D; Table S7).

### Association between the nucleotide and amino acid variation in *vrille* and variation in fly cold tolerance traits among clinal *D. montana* and *D. flavomontana* populations

To elucidate the adaptive potential of *vrille* in cold tolerance among *D. montana* and *D. flavomontana* populations, we traced the association between pairwise differences in *vrille* sequence variation (the number of nucleotide and amino acid differences) and differences in fly cold tolerance traits among the studied populations. The number of amino acid differences and the differences in mean CT_min_ in *D. montana* populations were significantly correlated (Mantel test: r=0.60, P=0.007; Fig. 5B; Table S8), i.e. the greater the amino acid differences were between *D. montana* populations, the greater were the differences in their chill coma resistance. Other measured correlations were non-significant (Fig. 5; Table S8).

**Figure 5.**
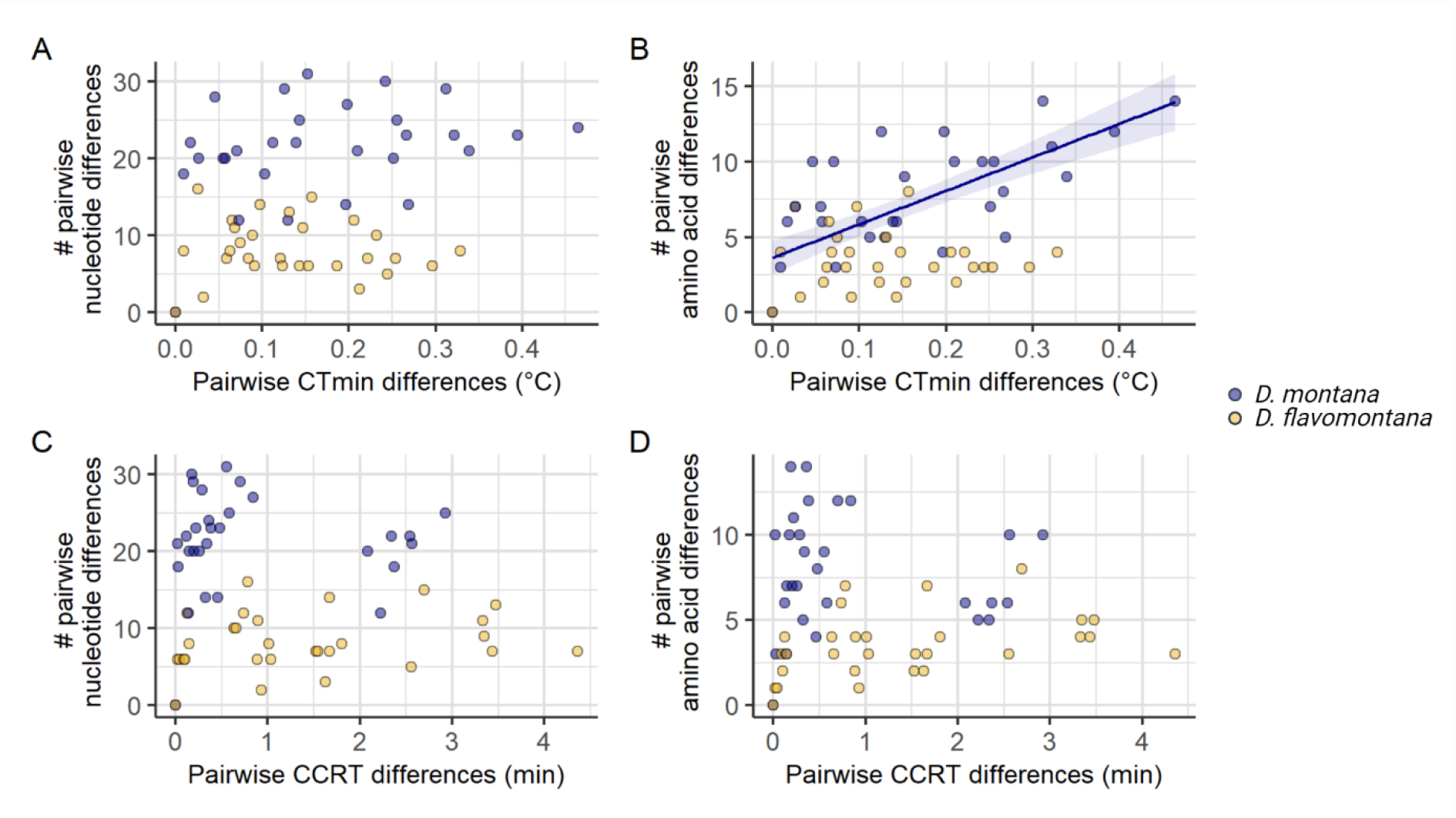
The association between *vrille* sequence variation and cold tolerance among *D. montana* and *D. flavomontana* populations. (A) Pairwise mean CT_min_ differences and the number of pairwise nucleotide differences among *D. montana* and *D. flavomontana* populations. (B) Pairwise mean CT_min_ differences and the number of pairwise amino acid differences among *D. montana* and *D. flavomontana* populations. (C) Pairwise mean CCRT differences and the number of pairwise nucleotide differences among *D. montana* and *D. flavomontana* populations. (D) Pairwise mean CCRT differences and the number of pairwise amino acid differences among *D. montana* and *D. flavomontana* populations

### Daily rhythm of *vrille* and the expression levels of *vrille* after RNAi in *D. montana*

Since the expression levels of circadian clock genes are known to fluctuate during the day, we first investigated daily rhythm in *vrille* expression in LD (light:dark cycle) 18:6, where the females of the study population of *D. montana* (Seward, Alaska, USA) can be expected to be non-diapausing (and which was also verified, see methods). Our results showed that *vrille* has a clear daily rhythm and that its highest expression corresponds to ZT16 and ZT20 (ZT = Zeitgeber Time which refers to the number of hours after the lights are switched on; see Fig. 6A). From these two time points we chose to perform RNAi-injections at ZT16, when the lights were still on.

**Figure 6.**
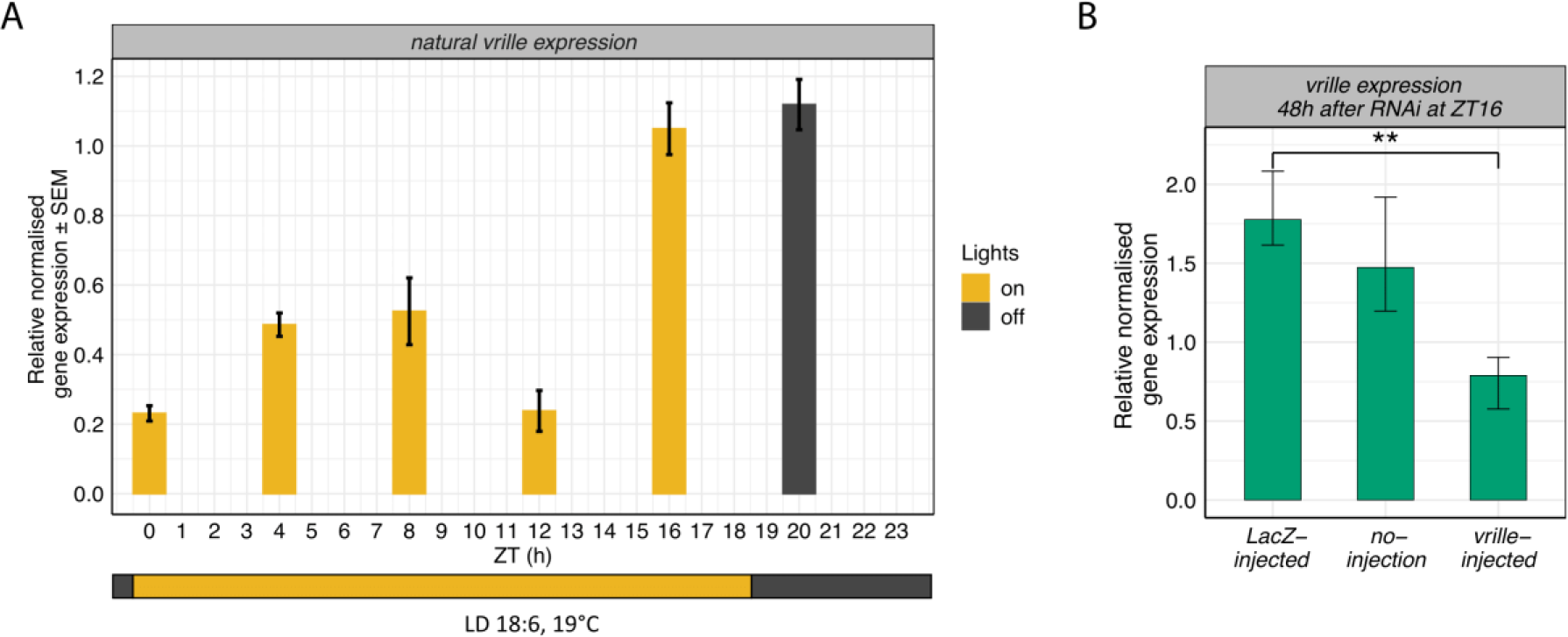
*vrille* expression during the day and after RNA interference (RNAi). (A) Relative normalised expression of naturally cycling *vrille* at six time points, starting at ZT0 (Zeitgeber Time = sampling time every 4 hours over a 24 hour period), in LD 18:6 and 19 °C. Yellow represents the time of the day when the lights were on and dark grey when the lights were off. (B) Relative normalised expression levels of *vrille* in *LacZ*-injected and no-injection control females, and in females injected with dsRNA targeting on *vrille* 48h after the injections at ZT16. Error bars represent bootstrapped 95% confidence intervals. Significance levels were obtained from linear model and only significant observations are shown: * P < 0.05, ** P < 0.01 and *** P < 0.001.

We next compared *vrille* expression levels in *LacZ*-injected control females to those of *vrille*-injected and no-injection females 12, 24 and 48 hours after the injections (Fig. S1, Fig. 6B). This enabled us to measure the effects of *vrille*-RNAi on the expression level of this gene controlling possible effects of immune responses and physical damage. Differences between *LacZ*- and *vrille*-injected females were most pronounced 48h after the RNAi-injections (Fig. 6B; Fig. S1), where *vrille*-injected females had approximately 56% lower *vrille* expression compared to *LacZ* controls (Fig. 6B; Table S9). Accordingly, all RNAi experiments were performed 48h after the injections at ZT16 in LD 18:6.

### The effects of *vrille*-RNAi on *D. montana* females’ cold tolerance and cold acclimation ability

The effects of *vrille*-RNAi on female cold tolerance and cold acclimation ability were studied by quantifying these traits in *vrille*-injected, *LacZ*-injected and no-injection females. Comparisons between *LacZ*- and *vrille*-injected females enabled us to determine whether a expression of *vrille* increases *D. montana* females’ ability to resist chill coma (CT_min_), to recover from it (CCRT) and/or to achieve higher cold-tolerance after cold acclimation. Comparisons between *LacZ*-injected and no-injection females, on the other hand, revealed possible immune responses to dsRNA and/or physical damage caused by the injection itself. All cold tolerance tests were started 48h after the injections at ZT16, when the effects of *vrille*-RNAi were at a highest level (Fig. 6, Fig. S1). Cold tolerance tests (CT_min_ and CCRT) were conducted the same way as for the females of cline populations, except that now the females were maintained in LD 18:6 instead of LL throughout the experiments (also during the acclimation treatment). Moreover, to measure females’ cold acclimation ability, half of the females (cold acclimation group) were maintained in 6 °C and the other half (non-acclimation group) in 19 °C for 5 days prior to cold tolerance tests. RNAi-injections were performed two days (48 h) before finishing acclimation treatment and performing the tests.

We first investigated whether cold acclimation in 6 °C (cold-acclimated females) had improved female cold tolerance compared to the flies kept in 19 °C (non-acclimated females) within the three experimental groups (*LacZ*-injected females, no-injection females and *vrille*-injected females). In CT_min_ tests, cold acclimation improved the chill coma resistance in *LacZ*-injected and no-injection females by decreasing their CT_min_ on average by 0.7 °C and 0.6 °C, respectively, while acclimation had no significant effect on the chill coma resistance of *vrille*-injected females (Fig. 7A; Table S10). In CCRT tests, cold-acclimated no-injection females recovered from chill coma on average 1.5 minutes faster than the non-acclimated ones (Fig. 7B; Table S10), i.e. their ability to recover from coma was faster, as could be expected. However, cold acclimation had no significant effect on the recovery time of *LacZ*-injected females, which suggests that either immune responses or physical damage had overridden the positive effects of the acclimation (Fig. 7B; Table S10). Finally, CCRT tests for *vrille*-injected females showed that cold acclimation had slowed down their recovery time by ∼3.5 minutes instead of fastening it (Fig. 7B; Table S10). Such a significant effect in females’ ability to recover from chill coma cannot be explained solely by immune responses or physical damage, which means that expression of *vrille* is essential for females’ cold acclimation ability.

**Figure 7.**
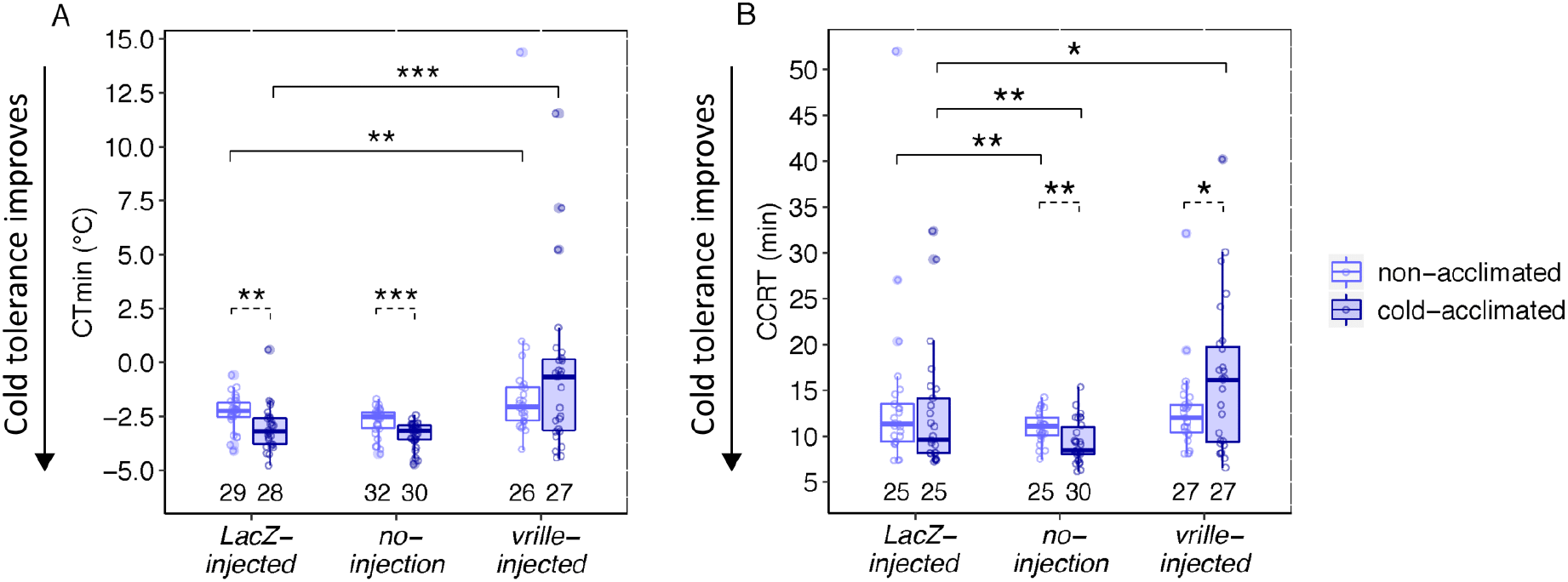
*D. montana* cold tolerance after RNA interference (RNAi). (A) Chill coma resistance (CT_min_) and (B) chill coma recovery time (CCRT) for *LacZ*-RNAi, no-injection, and *vrille*-RNAi females that were kept at +19 °C for the whole time, or at first at +19 °C and the last 5 days at +6 °C (cold acclimation period). Dashed lines indicate significant effects of the acclimation in each treatment, and solid lines significant differences between the *LacZ* control and the other treatments in females that were or were not acclimated. Significance levels were obtained from GLMMs and only significant observations are shown: * P < 0.05, ** P < 0.01 and *** P < 0.001. Numbers below boxplots refer to sample sizes. Whiskers represent ±1.5×IQR.

Next, we compared CT_min_ and CCRT of *LacZ*-injected females to those of the no-injection and *vrille-*injected females separately among the non-acclimated and cold-acclimated females. In CT_min_ tests, chill coma resistance did not differ significantly between *LacZ*-injected and no-injection females in either acclimation groups (Fig. 7A; Table S11). On the other hand, *vrille*-injected females entered chill coma in 1 °C (non-acclimated females) and 2.5 °C (cold-acclimated females) higher temperature than *LacZ*-injected females of the same groups (Fig. 7A; Table S11). These results show that low expression levels of *vrille* significantly decreased females’ ability to resist low temperatures. In CCRT tests, both the non-acclimated and cold-acclimated *LacZ*-injected females recovered from chill coma ∼3 minutes more slowly than the respective no-injection females (Fig. 7B; Table S11), which suggests that immune and/or injection effects might have played a role in chill coma recovery in both groups. The recovery time of non-acclimated *vrille*-injected females was on the same level as that of the *LacZ*-injected females (Fig. 7B; Table S11). However, cold-acclimated *vrille*-injected females recovered from chill coma ∼3.5 minutes more slowly than *LacZ*-injected females (Fig. 7B; Table S11), which again brings up the poor acclimation ability of *vrille*-injected females.

## Discussion

Numerous studies have found inter- and intraspecific latitudinal variation in insect cold tolerance (e.g. [1–4, 6, 16, 27–30]) and identified candidate genes for it (e.g. [62–64, 66]).

Spatially varying selection on insect fitness along latitudinal clines can be based on photoperiod and/or climatic factors, and sometimes it is difficult to detect the actual selection pressures driving the evolution of cold tolerance. To deepen our understanding on clinal adaptation, it is important to gather several independent sources of evidence, including sibling species, multiple populations or geographic regions and environmental correlations that account for population structure [79], as well as to investigate associations between cold tolerance and morphological traits (e.g. body colour and size) that often show parallel clines. Here, we explored factors affecting cold tolerance of non-diapausing females of *D. montana* and *D. flavomontana* originating from diverse climatic environments across different latitudes on the western coast and the Rocky Mountains of North America. Moreover, tracing sequence variation in both species and performing RNAi experiments in *D. montana* on one of the key regulators of the circadian behavioural rhythms, *vrille*, enabled us to investigate the role of this gene on female cold tolerance and cold acclimation ability.

### Effects of latitude and bioclimatic variables on fly cold tolerance vary even between closely-related species

The principal component analysis (PCA), which we performed on macroclimatic variables on fly collection sites, showed PC1 to separate the low-altitude coastal sites from the high-altitude mountainous ones based on differences in winter temperatures, temperature variability and precipitation. PC2 further arranged the clinal populations according to their summer and annual mean temperatures. Thus, in our dataset latitude represents photoperiodic differences (day length) between the sites, while PC2 corresponds to latitudinally varying mean temperatures and PC1 to macroclimatic variation between the coast and mountains.

Overall, genetic variation was lower in chill coma resistance (CT_min_) than in chill coma recovery time (CCRT), and CT_min_ showed significant variation only in *D. montana*. There are some plausible explanations for this pattern. CT_min_ may not be as suitable as CCRT for studying insects’ inherent cold tolerance, as rapid cold hardening (RCH) during the gradual cooling period in CT_min_ tests can change the composition of membrane phospholipids, improving the chill coma resistance [19, 20, 80, 81]. A trade-off between inherent cold tolerance and RCH, detected in several studies [26, 56, 82, 83], could lead to skewed or unapparent population-level variation in CT_min_. For example, sub-Antarctic *Ectemnorhinus* weevils and *Glossina pallidipes* tsetse flies show low inherent genetic variation and greater plastic responses in CT_min_ between populations [84, 85]. In contrast, CCRT has high heritability [27, 56, 86–88], and fast recovery from chill coma may be under strong selection due to energetically costly repair of cold injury and high fitness advantages involved in it (e.g. earlier recovery ensures high-quality territories, mates, feeding and breeding sites and escape from predators) [15, 18, 21–25]. These characteristics make CCRT a good indicator of climatic adaptation. Latitudinal variation in CCRT, but not in CT_min_, has also been observed in two widespread ant species, *Myrmica rubra* and *Myrmica ruginodis* [29]. Although it is possible that the low variation in cold tolerance in our study is a consequence of relatively low number of studied strains or populations, a recent study found similar trends when using mass-bred *D. montana* populations for a limited number of places from North America [89].

We predicted that cold tolerance traits have evolved in response to photoperiod, because it serves as a more reliable cue for seasonal temperature changes than environmental temperature itself [7]. In contrast to our expectations, *D. montana* CT_min_ was associated with latitudinally varying summer and annual mean temperatures (PC2) instead of photoperiod (latitude). This finding could indicate that the non-diapausing *D. montana* flies have adapted to cope with sudden temperature drops and diurnal temperature cycles at their home sites during the late spring, summer and early autumn [53]. CCRT of both species, on the other hand, was associated with photoperiod (latitude), as hypothesized. In *D. montana*, northern females recovered faster from chill coma than the southern ones independent of local climatic conditions (PC1 or PC2), while in *D. flavomontana* CCRT was also associated with macroclimatic conditions varying between the coastal and mountain sites (PC1). Surprisingly, *D. flavomontana* females from the colder high-altitude mountain populations recovered more slowly from chill coma than the ones from the humid low-altitude coastal populations. The most logical explanation for this is that high daily and seasonal temperature variation in the mountains create opposing selection pressures, which select for plastic responses. Accordingly, the flies could compensate their low inherent cold tolerance with high cold acclimation ability, as observed in several studies [26, 56, 82, 83, 90, 91]. For example, non-diapausing *D. montana* females from the coast have been shown to recover faster from chill coma than those from the mountains, but diapausing females from both regions recover at similar rates [91]. Mountain *D. flavomontana* females could also enhance their cold tolerance by entering reproductive diapause at earlier time of the year than females from the coastal populations on the same latitude, like *D. montana* females do [92]. Finally, the mountain flies could occupy lower mountain slopes, or thick snow cover could protect them during the cold season.

### Fly body colour and size and the effects of these traits on fly cold tolerance

Dark cuticle pigmentation (high melanism) can be expected to offer an advantage in cold environments, as it increases insects’ ability to absorb solar radiation and enables them to warm up faster in cold environments (thermal melanin hypothesis, [36]). This assumption has received support e.g. from latitudinal variation in the degree of melanism in 473 European butterfly and dragonfly species [33]. However, clinal variation in body colour can be induced also by selection favouring light individuals in the south due to their higher avoidance of UV radiation (UV protection hypothesis, [37, 38]). Overall, body colour genes are highly pleiotropic and can play a role also in a variety of other processes, including immunity, camouflage and mate choice [93, 94]. In our study, *D. montana* flies were darker and more cold-tolerant than *D. flavomontana* flies, which could be expected as *D. montana* is found on higher latitudes and altitudes than *D. flavomontana* [69, 70]. Body colour of *D. montana* showed no significant variation between populations, while that of *D. flavomontana* showed two trends. In the Rocky Mountains, *D. flavomontana* flies became darker towards North, as predicted by thermal melanin hypothesis [36], but the lack of association between fly colour and cold tolerance reduced the support to this theory (see [36]). Rocky Mountains cline could also be explained by UV protection hypothesis [37, 38], if the light flies collected from the southern populations proved to have higher UV resistance. The second trend, detected in *D. flavomontana* body colour, was increased melanism in the coastal populations. Also this trend could be explained by UV protection hypothesis, as the flies of the misty coastal populations receive less UV radiation than the ones living on high mountains. Desiccation resistance, linked with high melanism in many systems [39, 40, 95], is not likely to play an important role in the formation of either trend, as dark *D. flavomontana* individuals inhabit humid rather than arid regions. Sexual selection could play part in *D. flavomontana* colour variation, as cuticular hydrocarbons (CHCs), which can be regulated by pigmentation genes [96], are under sexual selection in this species [71]. Moreover, the darkness of *D. flavomontana* flies throughout the western coast could be explained by a founder effect, since this species has only recently distributed to the coastal area through British Columbia in Canada, where its body colour is dark [69, 71]. Finally, *D. flavomontana* may hybridise with *D. montana* to some degree [69–71], which could potentially have led to introgression of dark body colour from *D. montana* to *D. flavomontana*.

We hypothesized that body size (measured as weight) of our study species increases towards cold environments like e.g. in *Drosophila serrata* [27], consistent with Bergmann’s rule [45–47], and that the body size is correlated with cold tolerance traits. Contrary to our expectations, the body size of both species was largest in sites with warm summers and winters, which gives support for the converse Bergmann’s rule [45-47]. This likely results from the short growing season in cold regions and long generation time of *D. montana* and *D. flavomontana* (∼7 weeks) when the time available for development, growth and foraging is limited, resulting in smaller individuals [45-47]. However, *D. montana* body size is clearly affected by multiple selection pressures. While the body size of these flies decreased towards cold environments at population level, the large individuals recovered faster from chill coma than the smaller ones within populations. The latter finding is consistent with a previous study, where the overwinter survival of *D. montana* males was found to increase along with an increase in body size in nature [49]. In *D. flavomontana*, the largest individuals came from the southern Rocky Mountains and from the coastal region with warm summers and mild winters. Lack of correlation between body size and cold tolerance, detected in this species, resembles the situation in several other insect species [97-100].

### Circadian clock gene *vrille* plays an essential role in *D. montana* females’ short- and long-term cold acclimation

Insects’ circadian clock system monitors changes in daily light and temperature cycles and entrains behavioural and physiological rhythms to match with them [57]. This clock system shows adaptive divergence between northern and southern species [72-75], and several clock genes have also been found to be differentially expressed during cold acclimation in studied species [62, 63, 66], suggesting an association between the circadian clock system and cold adaptation. Here, we traced correlation between the number of nucleotide or amino acid differences in *vrille* and variation in the mean chill coma resistance (CT_min_) or chill coma recovery time (CCRT) in *D. montana* and *D. flavomontana* populations, and found a significant correlation only between amino acid variation and CT_min_ in *D. montana.* We then used RNAi to silence *vrille* and to investigate its effects on *D. montana* females’ cold tolerance traits and cold acclimation ability, and found this gene to play an important role in CT_min_ and in females’ cold acclimation ability, but not in CCRT. These findings together highlight the importance of *vrille* in enhancing females’ cold tolerance both during the rapid cold hardening occurring in CT_min_ test [81] and during the longer-term cold acclimation. Both traits, short- and long-term cold acclimation, have important ecological implications in cold environments with high daily and seasonal temperature changes.

Although we found a link between *vrille* and short- and long-term cold acclimation, it is not clear what are the genetic mechanisms underlying this link and to what extent the link exists. First, *vrille* might contribute to cold acclimation ability via circadian clock system, which may either directly contribute to cold acclimation, or regulate other pathways contributing to it. Here, any disruptions in the clock genes could cause similar effects. Alternatively, *vrille* might have more direct role on cold acclimation ability. For example, circadian clock system has been linked to Na^+^/K^+^ ion channel activity in *D. melanogaster* [101], which is also involved in avoiding cold injury, as observed in *D. montana* [102]. The second open question is whether the direct or indirect effects of *vrille* are limited to *D. montana*, *D. virilis* group species, or insects or organisms in general. *D. virilis* group species are among the most cold-tolerant *Drosophila* species [1, 2], their circadian clock system is adapted to northern habitats [72-75] and *vrille* expression is highly upregulated during cold-acclimation [64, 65], which gives indirect support for the role of *vrille* or the circadian clock system in cold adaptation in this group.

Although becoming more feasible, pinpointing the functions of genes affecting ecologically relevant traits is still rare. Previous RNAi experiments have verified e.g. *Hsp22 and Hsp23* genes to contribute to CCRT in *D. melanogaster* [12], *myo-inositol-1-sphosphate synthase* (*Inos*) to contribute to *D. montana* flies’ survival during cold stress (5 °C), but not to affect their CCRT [13], and *Gs1-like* (*Gs1l*) gene to be involved in *D. montana* cold acclimation [103]. *vrille* is an interesting addition to this list, and while the details on the specific molecular mechanisms remain unclear, it gives new insights on the genetics underlying insect cold tolerance and cold acclimation.

## Conclusions

Studying the mechanisms that generate and maintain variation in species stress tolerances is essential for understanding adaptation processes. We show that insect cold tolerance may rely on different environmental cues and morphological traits even in closely-related species, and that insects’ chill coma recovery time may be affected by stronger selection pressures in nature than chill coma resistance. These kinds of studies may have applications e.g. in predicting the likely outcomes of climate change or invasion biology. Species, whose cold tolerance is tightly linked with photoperiod, may encounter more difficulties in adapting to changing temperature conditions than the ones whose tolerances are driven by local climatic conditions. We also propose that *vrille*, and possibly the whole circadian clock system, play an essential role in molecular mechanisms underlying short- and long-term cold acclimation, both of which are ecologically important traits on high latitudes with high daily and seasonal temperature variation. In the future, studies on the relationship of reproductive stage and cold tolerance traits in individuals originating from diverse climatic conditions could deepen our understanding on adaptation to seasonally varying environments. Also, investigating the effects of silencing the other core circadian clock genes under cold environment and in other organisms would give valuable insights on the role of the circadian clock system on cold acclimation in general.

## Methods

### Genetic variation in cold tolerance, body colour and body size in *D. montana* and in *D. flavomontana* populations

#### Study species and populations

*D. montana* and *D. flavomontana* belong to the *montana* phylad of the *Drosophila virilis* group, and our recent whole genome analyses have shown their divergence time to be ∼1 mya (Poikela et al., in preparation). *D. montana* is distributed around the northern hemisphere across North America, Asia and Europe [70], while the distribution of *D. flavomontana* is restricted to North America [69, 70]. In the central Rocky Mountains, *D. montana* is found at altitudes from 1400 m to well over 3000 m, while *D. flavomontana* is found mainly below 2000 m. In the western coast, where *D. flavomontana* has probably invaded only during the last decades, both species live at much lower altitudes (see [69, 71]).

We investigated cold tolerance traits and measures of body colour and body weight using females from 23 *D. montana* and 20 *D. flavomontana* isofemale strains, which were established from the progenies of fertilized females collected from several sites in North America between 2013 and 2015 (Fig. 1). Each site was represented by three isofemale strains per population per species, when possible (Fig. 1; Table S1). *D. flavomontana* is particularly rare in the western coast of North America [71] and we succeeded to collect only 1-2 strains from the study populations. All the strains were maintained in continuous light at 19 ± 1 °C since their establishment (15-30 generations) to eliminate plastic responses and prevent females from entering reproductive diapause. For the experiments, we sexed emerging flies under light CO_2_ anaesthesia within three days after emergence. Females were changed into fresh malt-vials once a week and used in experiments at the age of 20 ± 2 days, when they all had fully developed ovaries [104].

#### Cold tolerance traits

We investigated variation in female cold tolerance using two well-defined and ecologically relevant methods: chill coma temperature (CT_min_; also called critical thermal minimum) and chill coma recovery time (CCRT). CT_min_ corresponds to the temperature, at which the fly resists cold until it loses all neurophysiological activity and coordination and falls into a chill coma during a gradual decrease in temperature (reviewed in [14]). In this test, we placed the females individually in glass vials, which were submerged into a 30 % glycol bath. We then decreased the bath temperature from the starting temperature of 19 °C at the rate of 0.5 °C/min and scored the CT_min_ for each fly. The second method, CCRT, measures the time taken from a fly to recover from a standardized exposure time at a chill-coma-inducing temperature (reviewed in [14]). In this test, we submerged the females individually in glass vials into a 30 % glycol bath for 16 hours at -6 °C [65]. After returning the vials into 19 ± 1 °C in fly laboratory, we measured the time required for each female to recover from the chill coma and stand on its legs. CT_min_ tests were replicated 21 times and CCRT tests 20 times with Julabo F32-HL Refrigerated/Heating Circulator and Haake k35 DC50 Refrig Circulating Bath Chiller, respectively. To account for possible variation between replicates, each test included 1-3 females from each strain.

#### Fly body colour and body weight

We analysed variation in the body colour and body weight (as a proxy of body size) of the same females that had been phenotyped in CT_min_ or CCRT tests. Immediately after the cold tolerance tests, the females were put individually into tightly sealed PCR plates and kept in -20 °C freezer until measuring their body colour and weight. For body colour measurements, the females were photographed under Leica L2 microscope with 5x magnification, using Leica DFC320 R2 camera together with the Leica Application Suite software v4.3.0. Exposure time, zoom factors and illumination level were kept constant, and the photographing arena was surrounded by a plastic cylinder to eliminate glares on the chitinous surface of the fly. All photographs were taken within 3 months after the cold tolerance tests. Images were saved in 24-bit (RGB) format and the colour intensity was measured using grayscale image analysis (ImageJ, [105]); linearly scaling from 0 to 255 (0 = black, 255 = white). We took colour measurements from part of thorax (scutum), as our preliminary tests showed that it best incorporates the colour variation among flies (Fig. S2). Body weight was measured by weighting the females with Mettler Toledo™ NewClassic Balance (model ME). The weight of the females used in CT_min_ tests was measured after females had been frozen for 2 to 4 months, which appeared to be problematic as the females had started to dry and their weight correlated with the freezing time (Fig. S3). The weight of the females used in CCRT tests was measured after 6 to 17 months in freezer. These females had lost most of their body liquids so that their weight was close to dry weight and freezing time had no effect on it (Fig. S3). Accordingly, only the latter dataset was used in the analyses.

#### Statistical analyses

We used different statistical models to investigate whether variation in fly cold tolerance, body colour and body weight was associated with latitude, as a proxy of photoperiod, and/or local climatic variables, and whether cold tolerance traits and morphological traits showed a correlation with each other. In these models, we used either CT_min_ data (chill coma temperatures in Celsius degrees + 10 °C to prevent negative values from affecting the analysis), CCRT data (in minutes), body colour (measured from CCRT flies) or body weight (mg; measured from CCRT flies) as response variables. *D. flavomontana* CT_min_ data were normally distributed (Fig. S4) and they were analysed with generalized linear mixed model (GLMM) with gaussian distribution (equivalent of linear mixed model). Other datasets showed deviation from the normality (Fig. S4), and they were analysed using generalized linear mixed model (GLMM) with gamma distribution, using *glmmTMB* function from glmmTMB package [106]. Technical replicates and isofemale strains were handled as crossed random effects.

In our dataset latitude and altitude were negatively correlated (Pearson correlation coefficient = -0.82) and to prevent this multicollinearity from affecting the analysis, we used only latitude as an explanatory factor. However, we considered the effect of altitude and climatic variables through a principal component analysis (PCA). We downloaded climatic information from WorldClim database v2.1 (2.5 min spatial resolution; current data 1970-2000; [107]; http://www.worldclim.org) and extracted 19 bioclimatic variables for each site using their latitudinal and longitudinal coordinates (Table S3; Fig. 1) with raster package v. 2.8-19 [108]. Moreover, we obtained growing season length (days) for each site from article by [92] and www.weatherbase.com. Growing season is defined as the average number of days per year when the average daily temperature is at least 5 °C. We performed the PCA on the 19 bioclimatic variables, altitude and growing season length (Table S3) to summarize climatic differences on temperature and precipitation in each site using “FactoMineR” package [109].

We included the first two PCs in the model comparison of cold tolerance, body colour and body weight (see Results). The simplest model included latitude and PC2, which enabled us to distinguish between the latitudinally varying photoperiods and temperatures. Moreover, the effect of macroclimatic conditions varying between the western coast and the Rocky Mountains (PC1), interaction terms between latitude/PC2 and PC1, and body colour (divided by 100 to scale the variables) and body weight were included in the model selection (Table S6). The best-fit model for CT_min_, CCRT, body colour and body weight of both species was chosen for further analysis based on Akaike information criterion (AIC) value and Akaike weight (Table S6) using *aictab* function from AICcmodavg package [110]. All the analyses were conducted in R (v1.2.1335-1) and R studio (v3.6.1).

### Investigating circadian clock gene *vrille* and its effects on fly cold tolerance using sequence variation and RNA interference (RNAi)

#### The candidate gene

*D. virilis* species group shows several distinct features in their circadian clock system compared to *D. melanogaster*, which has enabled their adaptation in high-latitudes [72-74]. Interestingly, *vrille* was the only core circadian clock gene that was found to show both population-level differentiation in clinal cold tolerance study [89] as well as expression level differences when cold-acclimated and non-acclimated *D. montana* flies were compared [77]. This gene was also one of the differentially expressed genes in our earlier microarray study investigating cold tolerance of *D. montana* and *D. virilis* [65]. Accordingly, we investigated the sequence variation in *vrille* and traced its association with cold tolerance traits (CT_min_ and CCRT) in *D. montana* and *D. flavomontana* populations. We also used RNAi to silence the expression of *vrille* and investigated its effects on cold tolerance traits (CT_min_ and CCRT) and cold acclimation ability in *D. montana*, where this gene showed correlation with CT_min_ at the amino acid level (see Fig. 5B).

#### Association between the number of nucleotide or amino acid differences and differences in cold tolerance traits in D. montana and D. flavomontana populations

To evaluate the role of *vrille* in cold adaptation, we investigated the association between *vrille* sequence variation and variation in cold tolerance traits (CT_min_ and CCRT) in *D. montana* and *D. flavomontana* populations. We obtained published *D. montana* genomic sequences from NCBI under accession number LUVX00000000 and the respective genome annotation from Dryad: doi:10.5061/dryad.s813p55 [111]. Based on *D. virilis* and *D. melanogaster* orthologs in the annotation file, *vrille* sequence is found in scaffold2327-size32305 at position 13933-17558. We used Illumina 150 bp paired-end data from single wild-caught females or their F1 female progeny, i.e. the founder females of the studied isofemale strains, from all study populations of both species (one individual per population; Table S1; Poikela et al., in prep.).

Before mapping, Illumina paired-end reads were quality-checked with FastQC v0.11.8 (Andrews 2010: FastQC: a quality control tool for high throughput sequence data. Available online at: http://www.bioinformatics.babraham.ac.uk/projects/fastqc) and trimmed for adapter contamination and low-quality bases using fastp v0.20.0 [112]. Each trimmed Illumina sample was mapped against *vrille D. montana* reference *vrille* gene using BWA mem v0.7.17 [113]. The alignments were sorted with SAMtools v1.10 [114] and PCR duplicates marked with sambamba v0.7.0 [115]. The resulting BAM-files were used for variant calling with freebayes v1.3.1-dirty [116], and the variant calling (VCF) file was filtered for minimum depth and quality of 20 using bcftools view v1.8 (SAMtools. 2018. https://github.com/samtools/bcftools/releases/tag/1.8). Finally, *vrille* FASTA sequences for each sample were obtained using bcftools consensus.

The *vrille* sequences together with information on exons from the annotation file (see Fig. S5, S6) were imported to Geneious v9.1.8. (Biomatters ltd). For exons, *D. montana* and *D. flavomontana* nucleotide sequences and translated amino acid sequences were aligned using muscle algorithm. The number of pairwise nucleotide and amino acid differences were then calculated for *D. montana* and *D. flavomontana* populations (Table S12, S13). Similarly, pairwise differences in mean CT_min_ (in Celsius degrees) and CCRT (in minutes) were calculated for *D. montana* and *D. flavomontana* populations (Table S14, S15). The correlations between distance matrices were statistically tested with a Mantel test with 1000 permutations using mantel.rtest function from ade4 package [117]. The correlations were tested between i) the number of nucleotide differences and differences in mean CT_min_ ii) the number of amino acid differences and differences in mean CT_min_, iii) the number of nucleotide differences and differences in mean CCRT, and iv) the number of amino acid differences and differences in mean CCRT among *D. montana* and *D. flavomontana* populations (only within-species correlations).

#### Study material for RNAi

We performed RNAi study using only the more cold-tolerant *D. montana*, where *vrille* showed correlation with CT_min_ at the amino acid level (see Fig. 5B) and which has become a model species for studying cold adaption at phenotypic and genetic level (e.g. [61, 64, 92]). Here, we used females of a mass-bred cage population originating from Seward (Alaska, USA; see Fig. 1) to increase genetic diversity. The population cage has been established from the 4th generation progenies of 20 isofemale strains (total of 400 flies) in 2013 and maintained in continuous light at 19 ± 1 °C since its establishment (30 generations). Malt bottles with freshly laid eggs were transferred from the population cage into a climate chamber in LD 18:6 (light:dark cycle) at 19 ± 1 °C to reinforce flies’ circadian rhythmicity prior to the experiments and ensure they are non-diapausing. Critical day length for the induction of reproductive diapause (CDL; 50 % of the females of given population enter diapause) in 19 °C is LD 17:7 in Seward population [92], and thus the females emerging in LD 18:6 can be expected to develop ovaries. After ∼4 weeks, we collected newly emerged females (≤ 1 day old) using light CO_2_ anaesthesia and placed them back into the above-mentioned conditions in malt-vials. Females were changed into fresh vials once a week until they were used in the experiments at the age of 21 days.

#### Defining daily expression rhythm of vrille for RNAi study

Expression levels of circadian clock genes are known to show daily fluctuations, and in order to perform RNAi experiments at a right time of the day, we defined the time when the expression of *vrille* is highest at LD 18:6. To do this, we collected females every four hours (i.e. at six time points / day), starting at 6 am (ZT0) when lights were switched on. At each time point, we stored samples of females into -80°C through liquid N_2_ and transferred them later on into RNAlater®-ICE frozen tissue transition solution (Thermo Fisher Scientific). We then checked the size of female ovaries (see [104]) and used only the females with fully developed ovaries (>95 % of the females). For each time point, RNA was extracted from three pools, each of which consisted of three whole females, using ZR Tissue & Insect RNA MicroPrep kit with DNAse treatment (Zymo Research^®^). RNA purity was analysed with NanoDrop® ND-1000 spectrophotometer (Thermo Fisher Scientific) and concentration with Qubit® 2.0 Fluorometer and RNA BR kit (Thermo Fisher Scientific). cDNA was synthesized using equal quantities of RNA (200 ng) with iScript Reverse Transcription kit (Bio-Rad Laboratories®).

We measured the expression levels of *vrille* with quantitative real time PCR (qPCR). qPCR primers for *vrille* and reference genes were designed based on *D. montana* genomic sequences under NCBI accession number LUVX00000000 [111], together with information from *D. virilis* exons using Primer3 (primer3.ut.ee) and NetPrimer (www.premierbiosoft.com/netprimer) programs (gene accession numbers and primer sequences in Table S16). qPCR mix contained 10 µl of 2x Power SYBR Green PCR Master Mix (Bio-Rad Laboratories), 0.3 µM of each gene-specific primer and 1 µl of cDNA solution. Cycling conditions in Bio-Rad CFX96 instrument were: 3 min at 95 °C, 10 sec at 95 °C, 10 sec at annealing temperature of 53 °C (for reference genes 56 °C) and 30 sec at 72 °C (repeated 40x), followed by melting curve analysis (65-95 °C) for amplification specificity checking. Each run included three technical replicates for each sample and the final threshold value (Cq) was defined as a mean of the technical replicates that produced good quality data. The relative qPCR data was normalised with ΔΔ(Ct) normalisation method [118] using two reference genes, *Tubulin beta chain* (*Tub2*) and *Ribosomal protein L32* (*RpL32*), that showed equal expression levels in all samples (data not shown). Real efficiency values of the genes used in the qPCR are given in Table S16.

#### Synthesis of double-stranded RNA for RNAi

*LacZ,* which codes a part of a bacterial gene, was used as a control for dsRNA injections. We generated fragments of *vrille* and *LacZ* genes, with the length of 347 and 529 bp, respectively, with PCR (primer information given in Table S16). PCR products were purified with GeneJET Gel Extraction kit (Thermo Fisher Scientific) and cloned using CloneJET PCR Cloning kit (Thermo Fisher Scientific). PCR products were ligated into the vector (pJET1.2/blunt Cloning vector), transformed into *E. coli* Zymo JM109 (Zymo Research) cells, which were grown on Luria Broth (LB) ampicillin plates. Individual colonies were picked up after 16 hours and cultivated overnight in LB solution with ampicillin using a Unimax 1010 shaker with incubator (Heidolph Instruments). The samples from the extracted bacterial solutions were analysed for the size of the products in the second PCR, which was carried out with pJET primers from the cloning kit, followed by agarose gel runs. We then selected the colonies with the right size products for the third PCR using pJET primers, where the R primer contained T7 promoter sequence at the 5’ end of the primer (primer sequences in Table S16). PCR products were first purified with GeneJet Gel Extraction kit and then used in transcription synthesis of the double-stranded RNA (dsRNA), using the TranscriptAid T7 High Yield Transcription kit (Thermo Fisher Scientific). Finally, we purified and precipitated the synthesized products with ethanol, suspended them in salt buffer and quantified them using NanoDrop and agarose gel.

#### RNAi microinjection procedure, response time screening and vrille expression at the chosen response time

Injecting dsRNA targeting *vrille* gene is expected to cause gene-specific effects, but it may also cause immune responses and physical damage (injection) in the flies. Accordingly, we used *LacZ* (encoding for bacterial gene) injections as a control for both immune response to dsRNA and to physical damage of injections, and no-injection as a baseline control. We checked the effectiveness of RNAi treatment on *vrille* expression 12h, 24h and 48h after injections, at a time of the day when it shows highest expression (ZT16, see Results). The females were injected into thorax with 138 nl of ∼20 µM dsRNA targeting *vrille* or *LacZ* using Nanoject II Auto-nanoliter Injector with 3.5” Drummond glass capillaries (Drummond Scientific) pulled with P-97 Flaming/Brown Micropipette Puller (Sutter Instrument). No-injection control females were not injected, but were otherwise handled in the same way as the injected females. To prevent CO_2_ anaesthesia from inducing variation between these groups, we injected six females at a time. For each response time (12h, 24h, 48h) all three treatments (*vrille*, *LacZ*, no-injection) were performed. Each treatment consisted of three pools, each pool containing 10 whole females. After reaching the response time, the females were transferred into -80 °C through liquid N_2_. Then, the females were transferred into RNAlater®-ICE solution and their ovaries were checked as explained above. RNA was extracted from the pools using TRIzol® Reagent with the PureLink® RNA Mini kit (Thermo Fisher Scientific). RNA purity was analysed with NanoDrop and concentration with Qubit and RNA BR kit. cDNA was synthesized using equal quantities of RNA (143 ng) using SuperScript IV First-Strand Synthesis System (Thermo Fisher Scientific). Expression levels of *vrille* 12, 24 and 48 hours after the injections were quantified with qPCR, as described above. The response time of 48 hours was the most effective (Fig. S1), and its effect was double-checked with another round of cDNA synthesis (200 ng) and qPCR (Fig. 6B).

#### Experimental design for studying the functional role of vrille in cold tolerance using RNAi

We investigated the role of *vrille* gene in resisting chill coma (CT_min_) and recovering from it (CCRT) in non-diapausing *D. montana* females originating from Seward, Alaska, USA. Moreover, we considered the plastic effects of *vrille* during cold acclimation. Prior to performing cold tolerance tests, females were maintained for 16 days in LD 18:6 at 19 °C, corresponding to summer conditions at their home site (LD 18:6 was used throughout the experiment). Then, half of these females were subjected to cold acclimation treatment at 6 °C for 3 days (cold-acclimated females; [65]), while the other half was maintained at 19 °C. At the age of 19 days, both cold-acclimated and non-acclimated females were collected from the chambers, anesthetized with CO_2_ and injected as described above. They were then placed back to either cold acclimation (6 °C) and non-acclimation (19 °C) conditions for two more days, as the expression levels of target genes had been found to be lowest 48h after RNAi treatment (Fig. S1). At the age of 21 days, females’ cold tolerance was quantified by measuring their chill coma temperature (CT_min_) or chill coma recovery time (CCRT) using Julabo F32-HL Refrigerated/Heating Circulator. Sample sizes for CT_min_ and CCRT tests were 26-32 and 25-30 females per treatment, respectively.

#### Statistical analyses of RNAi experiment

For investigating the effects of RNAi, expression levels of *vrille* in *LacZ*-injected females were compared to no-injection control females and females injected with dsRNA targeting on *vrille*. The relative normalised expression values were analysed using a linear model (ANOVA) in base R.

To test the cold acclimation effect within each treatment group (*vrille*-RNAi, *LacZ*-RNAi, no-injection control), we compared the cold tolerance of the females that had or had not been cold-acclimated. We then investigated the gene specific effects on cold tolerance by comparing *LacZ*-injected females to *vrille*-injected females and traced possible immune and physical effects of the injections by comparing *LacZ*-injected females to no-injection females. These analyses were performed separately for the cold-acclimated and non-acclimated females. All data showed deviation from normality (Fig. S7) and were analysed with generalized linear mixed model using gamma distribution (GLMM; *glmmTMB* function from glmmTMB package [106]. In the models, response variables were either CT_min_ (Celsius degrees + 10 to prevent negative values from affecting the results) or CCRT (minutes) data, and the test replicates were used as a random effect. All the analyses were conducted in R (v1.2.1335-1) and R studio (v3.6.1).

## Abbreviations

CT_min_: critical thermal minimum or chill coma temperature; cold tolerance method that measures the ability of an individual to resist cold and chill coma
CCRT: chill coma recovery time; cold tolerance method that measures the ability of an individual to recover from chill coma
RNAi: RNA interference; technique for gene silencing

## Declarations

### Ethics approval and consent to participate

Neither of the species is endangered, and the flies were collected along watersides on public lands outside National and State parks, where insect collecting does not require permits in the USA or Canada (The Wilderness Act of 1964, section 6302.15).

### Consent to publish

Not applicable.

### Availability of data and materials

The climatic data were obtained from WorldClim database v2.1, http://www.worldclim.org. The phenotypic data generated and analysed during the current study are available in the Dryad repository, https://doi.org/10.5061/dryad.98sf7m0fv. *D. montana* genome sequences were obtained from NCBI repository, under accession number LUVX00000000, and the respective genome annotation was obtained from the Dryad repository, https://doi.org/10.5061/dryad.s813p55. *vrille* sequence data (BAM files) analysed during the current study are available in the EMBL Nucleotide Sequence Database (ENA) repository, under accession number PRJEB45370.

### Authors’ Contributions

NP, VT, AH and MK designed the study. NP and MK performed the research. NP analysed the data and drafted the manuscript, and all authors finalised it.

## Acknowledgements

We thank Anna-Lotta Hiillos and Johanna Kinnunen for help in analysing fly photographs and performing cold tolerance experiments and Ville Hoikkala for help in finalizing the figures.

## References

1. Andersen JL, Manenti T, Sørensen JG, Macmillan HA, Loeschcke V, Overgaard J. How to assess *Drosophila* cold tolerance: chill coma temperature and lower lethal temperature are the best predictors of cold distribution limits. Funct Ecol. 2015;29:55–65.

2. Kellermann V, Loeschcke V, Hoffmann AA, Kristensen TN, Fløjgaard C, David JR, et al. Phylogenetic constraints in key functional traits behind species’ climate niches: patterns of desiccation and cold resistance across 95 *Drosophila* species. Evolution. 2012;66:3377–89.

3. Kimura MT. Cold and heat tolerance of drosophilid flies with reference to their latitudinal distributions. Oecologia. 2004;140:442–9.

4. Overgaard J, Kristensen TN, Mitchell KA, Hoffmann AA. Thermal tolerance in widespread and tropical *Drosophila* species: does phenotypic plasticity increase with latitude? Am Nat. 2011;178:80– 96.

5. Sunday JM, Bates AE, Dulvy NK. Global analysis of thermal tolerance and latitude in ectotherms. Proc R Soc B. 2011;278:1823–30.

6. Addo-Bediako A, Chown SL, Gaston KJ. Thermal tolerance, climatic variability and latitude. Proc R Soc B Biol Sci. 2000;267:739–45.

7. Angilletta MJ. Thermal adaptation: a theoretical and empirical synthesis. Oxford University Press.; 2009.

8. Doucet D, Walker VK, Qin W. The bugs that came in from the cold: molecular adaptations to low temperatures in insects. Cell Mol Life Sci. 2009;66:1404–18.

9. Helmuth B, Kingsolver JG, Carrington E. Biophysics, physiological ecology, and climate change: does melanism matter? Annu Rev Physiol. 2005;67:177–201.

10. Pörtner HO, Farrell AP. Physiology and climate change. Science (80-). 2008;322:690–2.

11. Somero GN. The physiology of climate change: how potentials for acclimatization and genetic adaptation will determine ‘winners’ and ‘losers.’ J Exp Biol. 2010;213:912–20.

12. Colinet H, Lee SF, Hoffmann A. Knocking down expression of Hsp22 and Hsp23 by RNA interference affects recovery from chill coma in *Drosophila melanogaster*. J Exp Biol. 2010;213:4146–50.

13. Vigoder FM, Parker DJ, Cook N, Tournière O, Sneddon T, Ritchie MG. Inducing cold-sensitivity in the frigophilic fly *Drosophila montana* by RNAi. PLoS One. 2016;11:1–9.

14. MacMillan HA, Sinclair BJ. Mechanisms underlying insect chill-coma. J Insect Physiol. 2011;57:12–20. doi:10.1016/j.jinsphys.2010.10.004.

15. Gibert P, Moreteau B, Pétavy G, Karan D, David JR. Chill-coma tolerance, a major climatic adaptation among *Drosophila* species. Evolution. 2001;55:1063–8.

16. Hoffmann AA, Anderson A, Hallas R. Opposing clines for high and low temperature resistance in *Drosophila melanogaster*. Ecol Lett. 2002;5:614–8.

17. Garcia MJ, Littler AS, Sriram A, Teets NM. Distinct cold tolerance traits independently vary across genotypes in *Drosophila melanogaster*. Evolution. 2020;74:1437–50.

18. Overgaard J, Macmillan HA. The integrative physiology of insect chill tolerance. Annu Rev Physiol. 2017;79:187–208.

19. Andersen JL, Macmillan HA, Overgaard J. Muscle membrane potential and insect chill coma. J Exp Biol. 2015;218:2492–5.

20. Macmillan HA, Findsen A, Pedersen TH, Overgaard J. Cold-induced depolarization of insect muscle: differing roles of extracellular K+ during acute and chronic chilling. J Exp Biol. 2014;217:2930–8.

21. Clark MS, Worland MR. How insects survive the cold: molecular mechanisms — a review. J Comp Physiol B. 2008;178:917–33.

22. MacMillan HA, Williams CM, Staples JF, Sinclair BJ. Reestablishment of ion homeostasis during chill-coma recovery in the cricket *Gryllus pennsylvanicus*. Proc Natl Acad Sci U S A. 2012;109:20750–20755.

23. Sinclair BJ, Gibbs AG, Roberts SP. Gene transcription during exposure to, and recovery from, cold and desiccation stress in *Drosophila melanogaster*. Insect Mol Biol. 2007;16:435–43.

24. Teets NM, Peyton JT, Ragland GJ, Colinet H, Renault D, Hahn DA, et al. Combined transcriptomic and metabolomic approach uncovers molecular mechanisms of cold tolerance in a temperate flesh fly. Physiol Genomics. 2012;44:764–77.

25. Colinet H, Lee SF, Hoffmann A. Temporal expression of heat shock genes during cold stress and recovery from chill coma in adult *Drosophila melanogaster*. FEBS J. 2010;277:174–85.

26. Hoffmann AA, Sørensen JG, Loeschcke V. Adaptation of *Drosophila* to temperature extremes: bringing together quantitative andmolecular approaches. J Therm Biol. 2003;28:175–216.

27. Hallas R, Schiffer M, Hoffmann AA. Clinal variation in *Drosophila serrata* for stress resistance and body size. Genet Res (Camb). 2002;79:141–8.

28. Overgaard J, Hoffmann AA, Kristensen TN. Assessing population and environmental effects on thermal resistance in *Drosophila melanogaster* using ecologically relevant assays. J Therm Biol. 2011;36:409–16. doi:10.1016/j.jtherbio.2011.07.005.

29. Maysov A. Chill coma temperatures appear similar along a latitudinal gradient, in contrast to divergent chill coma recovery times, in two widespread ant species. J Exp Biol. 2014;217:2650–8.

30. Castañeda LE, Lardies MA, Bozinovic F. Interpopulational variation in recovery time from chill coma along a geographic gradient: a study in the common woodlouse, Porcellio laevis. J Insect Physiol. 2005;51:1346–51.

31. Kozak KH, Graham CH, Wiens JJ. Integrating GIS-based environmental data into evolutionary biology. Trends Ecol Evol. 2008;23:141–8.

32. Hahn DA, Denlinger DL. Meeting the energetic demands of insect diapause: nutrient storage and utilization. J Insect Physiol. 2007;53:760–73.

33. Zeuss D, Brandl R, Brändle M, Rahbek C, Brunzel S. Global warming favours light-coloured insects in Europe. Nat Commun. 2014;5:1–9.

34. Heidrich L, Friess N, Fiedler K, Brändle M, Hausmann A, Brandl R, et al. The dark side of Lepidoptera: colour lightness of geometrid moths decreases with increasing latitude. Glob Ecol Biogeogr. 2018;27:407–16.

35. Clusella-Trullas S, Terblanche JS, Blackburn TM, Chown SL. Testing the thermal melanism hypothesis: a macrophysiological approach. Funct Ecol. 2008;22:232–8.

36. Clusella-Trullas S, van Wyk JH, Spotila JR. Thermal melanism in ectotherms. J Therm Biol. 2007;32:235–45.

37. Bastide H, Yassin A, Johanning EJ, Pool JE. Pigmentation in *Drosophila melanogaster* reaches its maximum in Ethiopia and correlates most strongly with ultra-violet radiation in sub-Saharan Africa. BMC Evol Biol. 2014;14:179.

38. Telonis-Scott M, Hoffmann AA, Sgrò CM. The molecular genetics of clinal variation: a case study of ebony and thoracic trident pigmentation in *Drosophila melanogaster* from eastern Australia. Mol Ecol. 2011;20:2100–10.

39. Rajpurohit S, Parkash R, Ramniwas S. Body melanization and its adaptive role in thermoregulation and tolerance against desiccating conditions in drosophilids. Entomol Res. 2008;38:49–60.

40. Ramniwas S, Kajla B, Dev K, Parkash R. Direct and correlated responses to laboratory selection for body melanisation in *Drosophila melanogaster*: support for the melanisation-desiccation resistance hypothesis. J Exp Biol. 2013;216:1244–54.

41. Kutch IC, Sevgili H, Wittman T, Fedorka KM. Thermoregulatory strategy may shape immune investment in *Drosophila melanogaster*. J Exp Biol. 2014;217:3664–9.

42. Chown SL, Gaston KJ. Body size variation in insects: a macroecological perspective. Biol Rev. 2010;85:139–69.

43. Klockmann M, Günter F, Fischer K. Heat resistance throughout ontogeny: body size constrains thermal tolerance. Glob Chang Biol. 2017;23:686–96.

44. Vinarski M V. On the applicability of Bergmann’s rule to ectotherms: the state of the art. Biol Bull Rev. 2014;4:232–42.

45. Blanckenhorn WU, Demont M. Bergmann and converse bergmann latitudinal clines in arthropods: two ends of a continuum? Integr Comp Biol. 2004;44:413–24.

46. Blanckenhorn WU, Tillwell RC, Young KA, Fox CW, Ashton KG. When Rensch meets Bergmann: does sexual size dimorphism change systematically with latitude? Evolution. 2006;60:2004–11.

47. Chown SL, Gaston KJ. Exploring links between physiology and ecology at macro-scales: the role of respiratory metabolism in insects. Biol Rev. 1999;74:87–120.

48. Hegna RH, Nokelainen O, Hegna JR, Mappes J. To quiver or to shiver: increased melanization benefits thermoregulation, but reduces warning signal efficacy in the wood tiger moth. Proc R Soc B Biol Sci. 2013;280:20122812.

49. Aspi J, Hoikkala A. Male mating success and survival in the field with respect to size and courtship song characters in *Drosophila littoralis* and *D. montana* (Diptera: Drosophilidae). J Insect Behav. 1995;8:67–87.

50. Franks SJ, Hoffmann AA. Genetics of climate change adaptation. Annu Rev Genet. 2012;46:185–208.

51. Kellermann V, van Heerwaarden B, Sgrò CM, Hoffmann AA. Fundamental evolutionary limits in ecological traits drive *Drosophila* species distributions. Science (80-). 2009;325:1244–7.

52. Vesala L, Salminen TS, Kostál V, Zahradníĉková H, Hoikkala A. Myo-inositol as a main metabolite in overwintering flies: seasonal metabolomic profiles and cold stress tolerance in a northern drosophilid fly. J Exp Biol. 2012;215:2891–7.

53. Teets NM, Denlinger DL. Physiological mechanisms of seasonal and rapid cold-hardening in insects. Physiol Entomol. 2013;38:105–16.

54. Denlinger DL. Relationship between cold hardiness and diapause. In: Insects at Low Temperature. Springer, Boston, MA; 1991. p. 174–98.

55. Vesala L, Salminen TS, Kankare M, Hoikkala A. Photoperiodic regulation of cold tolerance and expression levels of regucalcin gene in *Drosophila montana*. J Insect Physiol. 2012;58:704–9. doi:10.1016/j.jinsphys.2012.02.004.

56. Gerken AR, Eller OC, Hahn DA, Morgan TJ. Constraints, independence, and evolution of thermal plasticity: probing genetic architecture of long- and short-term thermal acclimation. Proc Natl Acad Sci. 2015;112:4399–404.

57. Barber AF, Sehgal A. Cold temperatures fire up circadian neurons. Cell Metab. 2018;27:951–3. doi:10.1016/j.cmet.2018.04.016.

58. Gunawardhana KL, Hardin PE. VRILLE controls PDF neuropeptide accumulation and arborization rhythms in small ventrolateral neurons to drive rhythmic behavior in *Drosophila*. Curr Biol. 2017;27:3442–53. doi:10.1016/j.cub.2017.10.010.

59. Hardin PE. Molecular genetic analysis of circadian timekeeping in *Drosophila*. Adv Genet. 2011;74:141–73.

60. Williams JA, Sehgal A. Molecular components of the circadian system in *Drosophila*. Annu Rev Physiol. 2001;63:729–55.

61. Kauranen H, Kinnunen J, Hiillos A, Lankinen P, Hopkins D, Wiberg RAW, et al. Selection for reproduction under short photoperiods changes diapause-associated traits and induces widespread genomic divergence. J Exp Biol. 2019;222:jeb205831.

62. Enriquez T, Colinet H. Cold acclimation triggers major transcriptional changes in *Drosophila suzukii*. BMC Genomics. 2019;20:1–17.

63. MacMillan HA, Knee JM, Dennis AB, Udaka H, Marshall KE, Merritt TJS, et al. Cold acclimation wholly reorganizes the *Drosophila melanogaster* transcriptome and metabolome. Sci Rep. 2016;6:1– 14. doi:10.1038/srep28999.

64. Parker DJ, Vesala L, Ritchie MG, Laiho A, Hoikkala A, Kankare M. How consistent are the transcriptome changes associated with cold acclimation in two species of the *Drosophila virilis* group? Heredity (Edinb). 2015;115:13–21.

65. Vesala L, Salminen TS, Laiho A, Hoikkala A, Kankare M. Cold tolerance and cold-induced modulation of gene expression in two *Drosophila virilis* group species with different distributions. Insect Mol Biol. 2012;21:107–18.

66. Des Marteaux LE, McKinnon AH, Udaka H, Toxopeus J, Sinclair BJ. Effects of cold-acclimation on gene expression in Fall field cricket (*Gryllus pennsylvanicus*) ionoregulatory tissues. BMC Genomics. 2017;18:1–17.

67. Espinoza C, Bieniawska Z, Hincha DK, Hannah MA. Interactions between the circadian clock and cold-response in *Arabidopsis*. Plant Signal Behav. 2008;3:593–4.

68. Gil KE, Park CM. Thermal adaptation and plasticity of the plant circadian clock. New Phytol. 2019;221:1215–29.

69. Patterson JT. Revision of the montana complex of the virilis species group. Univerisity Texas Publ. 1952;5204:20–34.

70. Throckmorton LH. The virilis species group. Genet Bioogy Drosoph. 1982;3:227–96.

71. Poikela N, Kinnunen J, Wurdack M, Kauranen H, Schmitt T, Kankare M, et al. Strength of sexual and postmating prezygotic barriers varies between sympatric populations with different histories and species abundances. Evolution. 2019;73:1182–99.

72. Menegazzi P, Benetta ED, Beauchamp M, Schlichting M, Steffan-dewenter I, Helfrich-Förster C. Adaptation of circadian neuronal network to photoperiod in high-latitude European Drosophilids. Cell. 2017;27:833–9.

73. Kauranen H, Ala-Honkola O, Kankare M, Hoikkala A. Circadian clock of *Drosophila montana* is adapted to high variation in summer day lengths and temperatures prevailing at high latitudes. J Insect Physiol. 2016;89:9–18. doi:10.1016/j.jinsphys.2016.03.005.

74. Kauranen H, Menegazzi P, Costa R, Helfrich-Förster C, Kankainen A, Hoikkala A. Flies in the North: locomotor behavior and clock neuron organization of *Drosophila montana*. J Biol Rhythms. 2012;27:377–87.

75. Bertolini E, Schubert FK, Zanini D, Sehadová H, Helfrich-Förster C, Menegazzi P. Life at high latitudes does not require circadian behavioral rhythmicity under constant darkness. Cell. 2019;29:3928–36.

76. Helfrich-Förster C, Bertolini E, Menegazzi P. Flies as models for circadian clock adaptation to environmental challenges. Eur J Neurosci. 2018;51:166–81.

77. Parker DJ, Envall T, Ritchie MG, Kankare M. Sex-specific responses to cold in a very cold-tolerant, northern *Drosophila* species. Heredity (Edinb). 2021;126:695–705. doi:10.1038/s41437-020-00398-2.

78. Aspi J, Lumme J, Hoikkala A, Heikkinen E. Reproductive ecology of the boreal riparian guild of *Drosophila*. Ecography (Cop). 1993;16:65–72.

79. Flatt T. Genomics of clinal variation in *Drosophila*: disentangling the interactions of selection and demography. Mol Ecol. 2016;25:1023–6.

80. Overgaard J, Sørensen JG, Petersen SO, Loeschcke V, Holmstrup M. Changes in membrane lipid composition following rapid cold hardening in *Drosophila melanogaster*. J Insect Physiol. 2005;51:1173–82.

81. Overgaard J, Sørensen JG, Petersen SO, Loeschcke V, Holmstrup M. Reorganization of membrane lipids during fast and slow cold hardening in *Drosophila melanogaster*. Physiol Entomol. 2006;31:328–35.

82. Noh S, Everman ER, Berger CM, Morgan TJ. Seasonal variation in basal and plastic cold tolerance: adaptation is influenced by both long- and short-term phenotypic plasticity. Ecol Evol. 2017;7:5248–57.

83. Nyamukondiwa C, Terblanche JS, Marshall KE, Sinclair BJ. Basal cold but not heat tolerance constrains plasticity among *Drosophila* species (Diptera: Drosophilidae). J Evol Biol. 2011;24:1927– 38.

84. Terblanche JS, Klok CJ, Krafsur ES, Chown SL. Phenotypic plasticity and geographic variation in thermal tolerance and water loss of the tsetse *Glossina pallidipes* (Diptera: Glossinidae): Implications for distribution modelling. Am J Trop Med Hyg. 2006;74:786–94.

85. Klok CJ, Chown SL. Resistance to temperature extremes in sub-Antarctic weevils: Interspecific variation, population differentiation and acclimation. Biol J Linn Soc. 2003;78:401–14.

86. Anderson AR, Hoffmann AA, McKechnie SW. Response to selection for rapid chill-coma recovery in *Drosophila melanogaster*: physiology and life-history traits. Genet Res. 2005;85:15–22.

87. Ayrinhac A, Gibert P, Legout H, Moreteau B, Vergilino R, David J. Cold adaptation in geographical populations of *Drosophila melanogaster*: phenotypic plasticity is more than genetic variability. Funct Ecol. 2004;18:700–6.

88. Bertoli CI, Scannapieco AC, Sambucetti P, Norry FM. Direct and correlated responses to chill-coma recovery selection in *Drosophila buzzatii*. Entomol Exp Appl. 2010;134:154–9.

89. Wiberg RAW, Tyukmaeva V, Hoikkala A, Ritchie MG, Kankare M. Cold adaptation drives population genomic divergence in the ecological specialist, Drosophila montana. bioRxiv. 2021.

90. Zuther E, Schulz E, Childs LH, Hincha DK. Clinal variation in the non-acclimated and cold-acclimated freezing tolerance of *Arabidopsis thaliana* accessions. Plant, Cell Environ. 2012;35:1860– 78.

91. Vesala L, Hoikkala A. Effects ofphotoperiodically induced reproductive diapause and cold hardening on the cold tolerance of *Drosophila montana*. J Insect Physiol. 2011;57:46–51.

92. Tyukmaeva VI, Lankinen P, Kinnunen J, Kauranen H, Hoikkala A. Latitudinal clines in the timing and temperature-sensitivity of photoperiodic reproductive diapause in *Drosophila montana*. Ecography (Cop). 2020;43:1–10.

93. Wittkopp PJ, Beldade P. Development and evolution of insect pigmentation: genetic mechanisms and the potential consequences of pleiotropy. Semin Cell Dev Biol. 2009;20:65–71.

94. True JR. Insect melanism: the molecules matter. Trends Ecol Evol. 2003;18:640–7.

95. Davis JS, Moyle LC. Desiccation resistance and pigmentation variation reflects bioclimatic differences in the *Drosophila americana* species complex. BMC Evol Biol. 2019;19:1–14.

96. Massey JH, Akiyama N, Bien T, Dreisewerd K, Wittkopp PJ, Yew JY, et al. Pleiotropic effects of ebony and tan on pigmentation and cuticular hydrocarbon composition in *Drosophila melanogaster*. Front Physiol. 2019;10:518.

97. Condon C, Acharya A, Adrian GJ, Hurliman AM, Malekooti D, Nguyen P, et al. Indirect selection of thermal tolerance during experimental evolution of *Drosophila melanogaster*. Ecol Evol. 2015;5:1873–80.

98. Gibert P, Huey RB. Chill-coma temperature in *Drosophila*: effects of developmental temperature, latitude, and phylogeny. Physiol Biochem Zool. 2001;74:429–34.

99. Norry FM, Loeschcke V. Temperature-induced shifts in associations of longevity with body size in *Drosophila melanogaster*. Evolution. 2002;56:299–306.

100. Scharf I, Sbilordo SH, Martin OY. Cold tolerance in flour beetle species differing in body size and selection temperature. Physiol Entomol. 2014;39:80–7.

101. Flourakis M, Kula-Eversole E, Hutchison AL, Han TH, Aranda K, Moose DL, et al. A conserved bicycle model for circadian clock control of membrane excitability. Cell. 2015;162:836–48. doi:10.1016/j.cell.2015.07.036.

102. MacMillan HA, Andersen JL, Davies SA, Overgaard J. The capacity to maintain ion and water homeostasis underlies interspecific variation in *Drosophila* cold tolerance. Sci Rep. 2015;5:1–11. doi:10.1038/srep18607.

103. Hopkins D, Envall T, Poikela N, Pentikäinen OT, Kankare M. Effects of cold acclimation and dsRNA injections on Gs1l gene splicing in *Drosophila montana*. Sci Rep. 2018;8:1–11.

104. Salminen TS, Hoikkala A. Effect of temperature on the duration of sensitive period and on the number of photoperiodic cycles required for the induction of reproductive diapause in *Drosophila montana*. J Insect Physiol. 2013;59:450–7. doi:10.1016/j.jinsphys.2013.02.005.

105. Schneider CA, Rasband WS, Eliceiri KW. NIH Image to ImageJ: 25 years of image analysis. Nat Methods. 2012;9:671–5. doi:10.1038/nmeth.2089.

106. Brooks ME, Kristensen K, van Benthem KJ, Magnusson A, Berg CW, Nielsen A, et al. glmmTMB balances speed and flexibility among packages for zero-inflated generalized linear mixed modeling. R J. 2017;9:378–400.

107. Fick SE, Hijmans RJ. WorldClim 2: new 1-km spatial resolution climate surfaces for global land areas. Int J Climatol. 2017;37:4302–15.

108. Hijmans RJ, Etten J Van. raster: Geographic data analysis and modeling. R package version 2.8–19. 2020.

109. Lê S, Josse J, Husson F. FactoMineR: an R Package for multivariate analysis. J Stat Softw. 2008;25:1–18.

110. Mazerolle M. AICcmodavg: model selection and multimodel inference based on QAICc. R package version 2.2–2, https://cran.r-project.org/package=AICcmodavg. 2019.

111. Parker DJ, Wiberg RAW, Trivedi U, Tyukmaeva VI, Gharbi K, Butlin RK, et al. Inter and intraspecific genomic divergence in *Drosophila montana* shows evidence for cold adaptation. Genome Biol Evol. 2018;10:2086–101.

112. Chen S, Zhou Y, Chen Y, Gu J. Fastp: An ultra-fast all-in-one FASTQ preprocessor. Bioinformatics. 2018;34:i884–90.

113. Li H, Durbin R. Fast and accurate short read alignment with Burrows – Wheeler transform. Bioinformatics. 2009;25:1754–60.

114. Li H, Handsaker B, Wysoker A, Fennell T, Ruan J, Homer N, et al. The Sequence Alignment / Map (SAM) Format and SAMtools. Bioinformatics. 2009;25:2078–2079.

115. Tarasov A, Vilella AJ, Cuppen E, Nijman IJ, Prins P. Sambamba: fast processing of NGS alignment formats. Bioi. 2015;31:2032–4.

116. Garrison E, Marth G. Haplotype-based variant detection from short-read sequencing. arXiv Prepr arXiv. 2012;:1207.3907 [q-bio.GN].

117. Dray S, Dufour AB. The ade4 package: implementing the duality diagram for ecologists. J Stat Softw. 2007;22:1–20.

118. Livak KJ, Schmittgen TD. Analysis of relative gene expression data using real-time quantitative PCR and the 2^-ΔΔC^T method. methods. 2001;408:402–8.

