## Supplementary information for "Multiple paths to cold tolerance: the role of environmental cues, morphological traits and the circadian clock gene *vrille*"

#### Content

Supplementary tables ............................................................................1
Supplementary figures ..........................................................................12
Supplementary references ……..............................................................18

#### Supplementary tables

Table S1. Information on fly collecting sites and years, and the exact coordinates (latitude, longitude) and altitudes for each collecting site. We collected flies of overwintered populations as soon as they started to fly (temperature increased above 12-15 ˚C). The collections were carried out in the northernmost populations in June and in the more southern ones in May. Table shows isofemale strains of both species (*mon* = *D. montana*, *fla* = *D. flavomontana*) used in the study. Single wild-caught females or their $F_1$ daughters, i.e. the founder females of the isofemale strains, were sequenced with Illumina (one individual per population and species; Poikela et al., in prep.).

| Site | Year | Latitude | Longitude | Altitude (m) | Species prefix and Strain ID | Illumina sample |
|---|---|---|---|---|---|---|
| Seward, AK, USA | 2013 | 60˚10’N | 149˚27’W | 35 | monSE13F14, -F16, -F37 | monSE13F37 |
| Terrace, BC, Canada | 2014 | 54˚27’N | 128˚34’W | 217 | monTER14F4, -F11, -F13<br>flaTER14F5 | monTER14F11<br>flaTER14F5 |
| McBride, BC, Canada | 2014 | 53˚07’N | 120˚18’W | 720 | monMB14F1, -F2, -F3<br>flaMB14F10, -F20, -F27 | monMB14F1<br>flaMB14F10 |
| Cranbrook, BC, Canada | 2014 | 49˚36’N | 115˚46’W | 940 | monCRAN14F16, -F20<br>flaCRAN14F7, -F10, -F13 | monCRAN14F16<br>flaCRAN14F7 |
| Vancouver, BC, Canada | 2014 | 49˚15’N | 123˚10’W | 4 | monVAN14F1, -F17, -F24<br>flaVAN14F9, -F20 | monVAN14F1<br>flaVAN14F20 |
| Ashford, WA, USA | 2013 | 46˚45’N | 121˚57’W | 573 | monASH13F9, -F11, -F13<br>flaASH13F2, -F14 | monASH13F13<br>flaASH13F2 |
| Livingston, MT, USA | 2013 | 45˚21’N | 110˚36’W | 1605 | flaMT13F8, -F11, -F15 | flaMT13F11 |
| Jackson, WY, USA | 2013 | 43˚26’N | 110˚50’W | 1857 | monJX13F3, -F41, -F48<br>flaJX13F31, -F37, -F38 | monJX13F48<br>flaJX13F31 |
| Afton, WY, USA | 2015 | 42˚43’N | 110˚55’W | 2000 | monAF15F12, -F19, -F28 | monAF15F28 |
| Liberty, UT, USA | 2015 | 41˚20’N | 111˚51’W | 1600 | flaLB15F1, -F2, -F3 | flaLB15F3 |

Table S2. A List of 19 bioclimatic variables used in the PCA (WorldClim database v2.1, 2.5 min spatial resolutions; current data 1970-2000; Fick and Hijmans 2017; www.worldclim.org).

| **Variable** | **Description** |
|---|---|
| bio1 | Annual mean temperature |
| bio2 | Mean diurnal range (mean of monthly (max - min temperature)) |
| bio3 | Isothermality (bio2/bio7*100) |
| bio4 | Temperature seasonality (standard deviation * 100) |
| bio5 | Max temperature of the warmest month |
| bio6 | Min temperature of the coldest month |
| bio7 | Annual temperature range (bio5-bio6) |
| bio8 | Mean temperature of the wettest quarter |
| bio9 | Mean temperature of the driest quarter |
| bio10 | Mean temperature of the warmest quarter |
| bio11 | Mean temperature of the coldest quarter |
| bio12 | Annual precipitation |
| bio13 | Precipitation of the wettest month |
| bio14 | Precipitation of the driest month |
| bio15 | Precipitation seasonality (coefficient of variation) |
| bio16 | Precipitation of the wettest quarter |
| bio17 | Precipitation of the driest quarter |
| bio18 | Precipitation of the warmest quarter |
| bio19 | Precipitation of the coldest quarter |

Table S3. 19 bioclimatic variables for each site were extracted from WorldClim database v2.1 using latitudinal and longitudinal coordinates (2.5 min spatial resolutions; current data 1970-2000; [1]; www.worldclim.org) and growing season lengths for each site were obtained from [2] and www.weatherbase.com.

| Population | longitude | latitude | altitude | bio1 | bio2 | bio3 | bio4 | bio5 | bio6 | bio7 | bio8 | bio9 | bio10 | bio11 | bio12 | bio13 | bio14 | bio15 | bio16 | bio17 | bio18 | bio19 | growing season length (days) |
|---|---|---|---|---|---|---|---|---|---|---|---|---|---|---|---|---|---|---|---|---|---|---|---|
| Seward | -149.45 | 60.16 | 35 | 3.84 | 6.65 | 28.52 | 643.90 | 16.80 | -6.54 | 23.34 | 3.56 | 10.33 | 12.13 | -3.44 | 1551 | 245 | 53 | 47 | 620 | 204 | 228 | 432 | 169 |
| Terrace | -128.57 | 54.45 | 217 | 6.41 | 7.72 | 27.57 | 730.96 | 21.38 | -6.63 | 28.01 | 1.75 | 13.43 | 15.32 | -2.71 | 1379 | 210 | 50 | 52 | 578 | 157 | 170 | 487 | 200 |
| McBride | -120.16 | 53.30 | 720 | 4.07 | 11.31 | 32.24 | 850.10 | 22.13 | -12.94 | 35.07 | 4.04 | 0.10 | 14.23 | -6.66 | 706 | 75 | 39 | 21 | 210 | 127 | 201 | 163 | 184 |
| Cranbrook | -115.74 | 49.52 | 940 | 5.55 | 12.24 | 32.16 | 899.03 | 25.95 | -12.10 | 38.06 | 14.11 | 0.93 | 16.50 | -5.85 | 461 | 61 | 26 | 27 | 151 | 85 | 133 | 113 | 188 |
| Vancouver | -122.60 | 49.24 | 4 | 10.04 | 9.15 | 36.73 | 577.06 | 24.20 | -0.70 | 24.90 | 3.52 | 17.02 | 17.09 | 3.06 | 1972 | 308 | 64 | 49 | 818 | 221 | 226 | 691 | 289 |
| Ashford | -121.96 | 46.76 | 573 | 6.25 | 9.40 | 39.81 | 512.91 | 20.72 | -2.88 | 23.60 | 1.02 | 12.65 | 13.00 | 0.69 | 2333 | 361 | 48 | 58 | 1057 | 199 | 213 | 964 | 227 |
| Livingston | -110.61 | 45.36 | 1605 | 5.13 | 14.06 | 38.05 | 836.35 | 25.90 | -11.05 | 36.95 | 12.93 | -4.85 | 15.66 | -4.85 | 500 | 72 | 24 | 35 | 185 | 83 | 154 | 83 | 193 |
| Jackson | -110.84 | 43.43 | 1857 | 3.51 | 15.73 | 38.05 | 924.38 | 26.55 | -14.80 | 41.35 | 8.07 | -1.79 | 14.87 | -8.04 | 456 | 52 | 28 | 20 | 132 | 94 | 110 | 120 | 173 |
| Afton | -110.92 | 42.72 | 2000 | 3.14 | 16.64 | 36.87 | 1030.70 | 27.49 | -17.63 | 45.12 | 8.47 | 15.39 | 15.39 | -10.24 | 466 | 53 | 30 | 16 | 140 | 104 | 104 | 112 | 176 |
| Liberty | -111.89 | 41.33 | 1600 | 5.95 | 13.73 | 36.51 | 866.64 | 27.02 | -10.59 | 37.61 | 4.65 | 16.83 | 16.83 | -4.49 | 515 | 55 | 27 | 21 | 162 | 89 | 89 | 129 | 239 |

Table S4. Principal components with their variance, cumulative variance and Eigenvalues.

| PC | Eigenvalue | Variance (%) | Cumulative variance (%) |
|---|---|---|---|
| PC1 | 14.39 | 68.52 | 68.52 |
| PC2 | 3.16 | 15.07 | 83.59 |
| PC3 | 1.39 | 6.60 | 90.18 |
| PC4 | 1.15 | 5.48 | 95.67 |
| PC5 | 0.39 | 1.86 | 97.53 |
| PC6 | 0.28 | 1.34 | 98.87 |
| PC7 | 0.14 | 0.66 | 99.53 |
| PC8 | 0.06 | 0.30 | 99.83 |
| PC9 | 0.04 | 0.17 | 100.00 |

Table S5. Contributions (loadings) of the altitude and 19 bioclimatic variables on the Principal Component (PC).

| Variable | PC1 | PC2 | PC3 | PC4 | PC5 |
|---|---|---|---|---|---|
| altitude | 5.5 | 1.3 | 10.5 | 0.2 | 1.6 |
| bio1 | 2.8 | 15.1 | 7.0 | 1.4 | 0.1 |
| bio2 | 5.5 | 3.0 | 7.4 | 0.0 | 1.8 |
| bio3 | 0.3 | 11.9 | 36.9 | 4.4 | 4.2 |
| bio4 | 6.5 | 0.1 | 0.8 | 2.7 | 0.3 |
| bio5 | 3.9 | 13.2 | 0.1 | 0.0 | 0.0 |
| bio6 | 6.3 | 1.7 | 0.4 | 1.8 | 1.4 |
| bio7 | 6.6 | 0.5 | 0.3 | 0.8 | 0.6 |
| bio8 | 3.7 | 0.3 | 1.9 | 23.8 | 5.4 |
| bio9 | 1.7 | 3.8 | 0.0 | 49.3 | 5.8 |
| bio10 | 0.8 | 20.4 | 16.1 | 0.0 | 0.0 |
| bio11 | 5.6 | 4.3 | 0.3 | 3.1 | 0.3 |
| bio12 | 6.6 | 0.1 | 3.2 | 0.0 | 0.2 |
| bio13 | 6.6 | 0.1 | 2.7 | 0.0 | 1.2 |
| bio14 | 6.0 | 0.0 | 1.0 | 2.6 | 13.1 |
| bio15 | 5.9 | 0.0 | 0.0 | 3.0 | 23.0 |
| bio16 | 6.4 | 0.1 | 4.1 | 0.0 | 2.0 |
| bio17 | 6.3 | 0.2 | 0.4 | 1.2 | 7.2 |
| bio18 | 5.1 | 2.5 | 0.1 | 5.5 | 24.8 |
| bio19 | 6.0 | 0.5 | 6.0 | 0.0 | 2.2 |
| growing season length | 2.0 | 20.8 | 0.7 | 0.1 | 4.7 |

Table S6. The best-fit model for CCRT, $CT_{min}$, body colour and size of *D. montana* and *D. flavomontana* was defined based on Akaike Information Criterion (AIC). The model selection included Latitude and PC2, as well as PC1, different interaction terms and body colour and size. The model with the highest Akaike weight (AICcWt), i.e. probability of being the best model, was chosen for further analysis. AICcWt of the best-fit model is shown in grey.

| **Model selection** | **$CT_{min}$** | | | | **$CT_{min}$** | | | | **CCRT** | | | | **CCRT** | | | |
|---|---|---|---|---|---|---|---|---|---|---|---|---|---|---|---|---|
| | ***D. montana*** | | | | ***D. flavomontana*** | | | | ***D. montana*** | | | | ***D. flavomontana*** | | | |
| **Model** | **df** | **AICc** | **ΔAICc** | **AICcWt** | **df** | **AICc** | **ΔAICc** | **AICcWt** | **df** | **AICc** | **ΔAICc** | **AICcWt** | **df** | **AICc** | **ΔAICc** | **AICcWt** |
| latitude + PC2 | 6 | 1046.1 | 0.0 | 0.4 | 6 | 979.8 | 0.4 | 0.3 | 6 | 2248.4 | 7.8 | 0.0 | 6 | 2238.7 | 0.7 | 0.7 |
| latitude + PC2 + PC1 | 7 | 1047.6 | 1.5 | 0.2 | 7 | 981.2 | 1.9 | 0.1 | 7 | 2249.0 | 8.4 | 0.0 | 7 | 2240.7 | 2.6 | 0.3 |
| latitude * PC1 + PC2 | NA | NA | NA | NA | NA | NA | NA | NA | 8 | 2250.4 | 9.8 | 0.0 | 8 | 2238.0 | 0.0 | 1.0 |
| latitude + PC2 * PC1 | 8 | 1049.3 | 3.2 | 0.1 | 8 | 983.3 | 3.9 | 0.1 | 8 | 2250.3 | 9.7 | 0.0 | 8 | 2242.7 | 4.7 | 0.1 |
| latitude * PC1 + PC2 * PC1 | NA | NA | NA | NA | NA | NA | NA | NA | 9 | 2252.2 | 11.6 | 0.0 | 9 | 2240.0 | 2.0 | 0.4 |
| latitude + PC2 + weight | | | | | | | | | 7 | 2240.6 | 0.0 | 1.0 | 7 | 2239.8 | 1.8 | 0.4 |
| latitude + PC2 + PC1 + weight | | | | | | | | | 8 | 2241.1 | 0.5 | 0.8 | 8 | 2241.7 | 3.7 | 0.2 |
| latitude * PC1 + PC2 + weight | | | | | | | | | 9 | 2242.2 | 1.6 | 0.4 | 9 | 2238.9 | 0.8 | 0.7 |
| latitude + PC2 * PC1 + weight | | | | | | | | | 9 | 2242.0 | 1.5 | 0.5 | 9 | 2243.8 | 5.7 | 0.1 |
| latitude * PC1 + PC2 * PC1 + weight | | | | | | | | | 10 | 2243.9 | 3.3 | 0.2 | 10 | 2241.0 | 2.9 | 0.2 |
| latitude + PC2 + colour | 7 | 1047.6 | 1.4 | 0.2 | 7 | 979.4 | 0.0 | 0.3 | 7 | 2249.2 | 8.6 | 0.0 | 7 | 2240.7 | 2.7 | 0.3 |
| latitude + PC2 + PC1 + colour | 8 | 1049.0 | 2.9 | 0.1 | 8 | 981.1 | 1.8 | 0.1 | 8 | 2249.5 | 8.9 | 0.0 | 8 | 2242.6 | 4.6 | 0.1 |
| latitude * PC1 + PC2 + colour | NA | NA | NA | NA | NA | NA | NA | NA | 9 | 2250.7 | 10.1 | 0.0 | 9 | 2239.6 | 1.6 | 0.5 |
| latitude + PC2 * PC1 + colour | 9 | 1050.6 | 4.5 | 0.0 | 9 | 983.1 | 3.7 | 0.1 | 9 | 2250.8 | 10.2 | 0.0 | 9 | 2244.7 | 6.7 | 0.0 |
| latitude * PC1 + PC2 * PC1 + colour | NA | NA | NA | NA | NA | NA | NA | NA | 10 | 2252.6 | 12.0 | 0.0 | 10 | 2241.7 | 3.7 | 0.2 |
| latitude + PC2 + weight + colour | | | | | | | | | 8 | 2242.3 | 1.7 | 0.4 | 8 | 2241.9 | 3.8 | 0.1 |
| latitude + PC2 + PC1 + weight + colour | | | | | | | | | 9 | 2242.6 | 2.0 | 0.4 | 9 | 2243.7 | 5.7 | 0.1 |
| latitude * PC1 + PC2 + weight + colour | | | | | | | | | 10 | 2243.6 | 3.0 | 0.2 | 10 | 2240.6 | 2.6 | 0.3 |
| latitude + PC2 * PC1 + weight + colour | | | | | | | | | 10 | 2243.6 | 3.0 | 0.2 | 10 | 2245.8 | 7.8 | 0.0 |
| latitude * PC1 + PC2 * PC1 + weight + colour | | | | | | | | | 11 | 2245.4 | 4.8 | 0.1 | 11 | 2242.7 | 4.7 | 0.1 |

| **Model selection** | **Body colour** | | | | **Body colour** | | | | **Body weight** | | | | **Body weight** | | | |
|---|---|---|---|---|---|---|---|---|---|---|---|---|---|---|---|---|
| | ***D. montana*** | | | | ***D. flavomontana*** | | | | ***D. montana*** | | | | ***D. flavomontana*** | | | |
| **Model** | **df** | **AICc** | **ΔAICc** | **AICcWt** | **df** | **AICc** | **ΔAICc** | **AICcWt** | **df** | **AICc** | **ΔAICc** | **AICcWt** | **df** | **AICc** | **ΔAICc** | **AICcWt** |
| latitude + PC2 | 6 | 2370.3 | 1.0 | 0.2 | 6 | 2301.7 | 9.4 | 0.0 | 6 | -129.7 | 0.0 | 1.0 | 6 | -118.0 | 5.3 | 0.1 |
| latitude + PC2 + PC1 | 7 | 2369.2 | 0.0 | 0.3 | 7 | 2297.4 | 5.1 | 0.1 | 7 | -127.7 | 2.0 | 0.4 | 7 | -119.9 | 3.4 | 0.2 |
| latitude * PC1 + PC2 | 8 | 2369.9 | 0.7 | 0.2 | 8 | 2295.2 | 2.9 | 0.2 | 8 | -125.6 | 4.1 | 0.1 | 8 | -117.8 | 5.5 | 0.1 |
| latitude + PC2 * PC1 | 8 | 2371.1 | 1.8 | 0.1 | 8 | 2292.3 | 0.0 | 0.8 | 8 | -126.0 | 3.7 | 0.2 | 8 | -123.3 | 0.0 | 1.0 |
| latitude * PC1 + PC2 * PC1 | 9 | 2370.3 | 1.0 | 0.2 | 9 | NA | NA | NA | 9 | -123.9 | 5.8 | 0.1 | 9 | -121.2 | 2.1 | 0.4 |

df = degrees of freedom
AICc = Akaike Information Criterion with a correction for small sample sizes
ΔAICc = difference between the best and other models
NA = model overparameterized given the data

Table S7. Summary of the best-fit model results on the effects of latitude and/or climatic factors (PC1) on cold tolerance and body colour of *D. flavomontana*. Model selection was based on Akaike Information Criterion (AIC) results (shown in Table S6). Significant P-values are shown in bold. df = degrees of freedom

| Species | Test | Fixed effect | Df | Estimate | SE | Statistic | P-value |
|---|---|---|---|---|---|---|---|
| ***D. montana*** | **$CT_{min}$** | Intercept | 7,443 | 2.023 | 0.059 | 34.42 | < 0.001 |
| | | Latitude | | 0.001 | 0.001 | 0.98 | 0.329 |
| | | PC2 | | 0.008 | 0.004 | 2.36 | **0.018** |
| ***D. flavomontana*** | **$CT_{min}$** | Intercept | 7,360 | 2.495 | 0.156 | 15.95 | < 0.001 |
| | | Latitude | | -0.002 | 0.002 | -1.00 | 0.316 |
| | | PC2 | | -0.005 | 0.006 | -0.95 | 0.341 |
| | | colour | | -0.123 | 0.078 | -1.58 | 0.114 |
| ***D. montana*** | **CCRT** | Intercept | 7,443 | 3.123 | 0.236 | 13.22 | < 0.001 |
| | | Latitude | | -0.011 | 0.005 | -2.48 | **0.013** |
| | | PC2 | | -0.009 | 0.014 | -0.61 | 0.540 |
| | | weight | | -0.210 | 0.069 | -3.07 | **0.002** |
| ***D. flavomontana*** | **CCRT** | Intercept | 7,385 | 3.729 | 0.357 | 10.44 | < 0.001 |
| | | Latitude | | -0.019 | 0.007 | -2.65 | **0.008** |
| | | PC1 | | 0.267 | 0.128 | 2.09 | **0.036** |
| | | Latitude × PC1 | | -0.006 | 0.003 | -2.07 | **0.038** |
| | | PC2 | | -0.013 | 0.019 | -0.70 | 0.484 |
| ***D. montana*** | **Body colour** | Intercept | 7,440 | 4.771 | 0.039 | 122.08 | < 0.001 |
| | | Latitude | | -0.001 | 0.001 | -1.60 | 0.109 |
| | | PC2 | | -0.002 | 0.002 | -0.96 | 0.339 |
| | | PC1 | | 0.002 | 0.001 | 1.83 | 0.067 |
| ***D. flavomontana*** | **Body colour** | Intercept | 7,384 | 5.080 | 0.069 | 73.79 | < 0.001 |
| | | Latitude | | -0.005 | 0.001 | -3.51 | **< 0.001** |
| | | PC2 | | -0.005 | 0.004 | -1.33 | 0.183 |
| | | PC1 | | -0.006 | 0.001 | -4.00 | **< 0.001** |
| | | PC1 × PC2 | | 0.002 | 0.001 | 2.94 | **0.003** |
| ***D. montana*** | **Body weight** | Intercept | 7,440 | -0.299 | 0.141 | -2.13 | 0.034 |
| | | Latitude | | 0.005 | 0.003 | 1.86 | 0.063 |
| | | PC2 | | 0.033 | 0.009 | 3.83 | **< 0.001** |
| ***D. flavomontana*** | **Body weight** | Intercept | 7,384 | -0.224 | 0.311 | -0.72 | 0.471 |
| | | Latitude | | 0.006 | 0.006 | 0.89 | 0.373 |
| | | PC2 | | 0.031 | 0.018 | 1.71 | 0.087 |
| | | PC1 | | 0.020 | 0.007 | 3.05 | **0.002** |
| | | PC1 × PC2 | | -0.010 | 0.004 | -2.60 | **0.009** |

Table S8. Summary of the correlations between two distance matrices, i.e. population differences in the mean $CT_{min}$ or CCRT and differences in the number of nucleotides or amino acids of *vrille* in *D. montana* and *D. flavomontana*, using a Mantel test with 1000 permutations.

| | Correlation | Observed correlation (r) | P |
|---|---|---|---|
| *D. montana* | $CT_{min}$ vs nucleotide differences | 0.21 | 0.178 |
| | $CT_{min}$ vs amino acid differences | 0.60 | **0.007** |
| | CCRT vs nucleotide differences | -0.11 | 0.533 |
| | CCRT vs amino acid differences | -0.14 | 0.542 |
| *D. flavomontana* | $CT_{min}$ vs nucleotide differences | -0.21 | 0.832 |
| | $CT_{min}$ vs amino acid differences | -0.12 | 0.676 |
| | CCRT vs nucleotide differences | 0.14 | 0.309 |
| | CCRT vs amino acid differences | 0.32 | 0.171 |

Table S9. Summary of the effect of treatment (*LacZ* and no-injection controls and RNAi with *vrille*) on the expression levels of *vrille*. Significant P-values are in bold.

| Gene | Treatment | Coefficient | SE | t | P |
|---|---|---|---|---|---|
| *vrille* | *LacZ* (intercept) | 1.777 | 0.169 | 10.538 | < 0.001 |
| | No-injection | -0.303 | 0.239 | -1.272 | 0.251 |
| | *vrille* | -0.988 | 0.239 | -4.143 | **0.006** |

Table S10. Summary of the effects of cold acclimation treatment on cold tolerance, measured with $CT_{min}$ or CCRT, in *D. montana* females in different treatments (*LacZ* and no-injection controls and RNAi with *vrille*) using generalized linear mixed model (GLMM) with gamma distribution. Significant P-values are in bold.

| Test | Acclimation effect | Treatment | Coefficient | SE | z-statistic | P-value | DF |
|---|---|---|---|---|---|---|---|
| **$CT_{min}$** | *LacZ*-injected | Intercept | 2.041 | 0.023 | 90.040 | < 0.001 | 1,53 |
| | | Acclimation: yes | -0.100 | 0.032 | -3.090 | **0.002** | |
| | No-injection | Intercept | 1.982 | 0.018 | 110.030 | < 0.001 | 1,58 |
| | | Acclimation: yes | -0.084 | 0.020 | -4.140 | **< 0.001** | |
| | *vrille*- injected | Intercept | 2.184 | 0.072 | 30.456 | < 0.001 | 1,49 |
| | | Acclimation: yes | 0.043 | 0.079 | 0.545 | 0.586 | |
| **CCRT** | *LacZ*- injected | Intercept | 2.611 | 0.115 | 22.756 | < 0.001 | 1,46 |
| | | Acclimation: yes | -0.089 | 0.116 | -0.764 | 0.445 | |
| | No-injection | Intercept | 2.386 | 0.046 | 51.480 | < 0.001 | 1,51 |
| | | Acclimation: yes | -0.154 | 0.054 | -2.880 | **0.004** | |
| | *vrille*- injected | Intercept | 2.547 | 0.076 | 33.520 | < 0.001 | 1,50 |
| | | Acclimation: yes | 0.237 | 0.107 | 2.200 | **0.028** | |

Table S11. Summary of the effects of silencing *vrille* gene on cold tolerance, measured with $CT_{min}$ or CCRT, in non-acclimated and cold-acclimated *D. montana* females using generalized linear mixed model (GLMM) with gamma distribution.

| Test | Gene effect | Treatment | Coefficient | SE | z | P | Df |
|---|---|---|---|---|---|---|---|
| **$CT_{min}$** | Non-acclimated | *LacZ*-injected (intercept) | 2.042 | 0.036 | 57.510 | < 0.001 | 2,80 |
| | | No-injection | -0.059 | 0.044 | -1.330 | 0.185 | |
| | | *vrille*- injected | 0.123 | 0.046 | 2.670 | **0.008** | |
| | Cold-acclimated | *LacZ*- injected (intercept) | 1.941 | 0.040 | 48.640 | < 0.001 | 2,82 |
| | | No-injection | -0.043 | 0.055 | -0.780 | 0.436 | |
| | | *vrille*- injected | 0.300 | 0.058 | 5.170 | **< 0.001** | |
| **CCRT** | Non-acclimated | *LacZ*- injected (intercept) | 2.597 | 0.087 | 29.940 | < 0.001 | 2,72 |
| | | No-injection | -0.248 | 0.085 | -2.918 | **0.004** | |
| | | *vrille*- injected | -0.054 | 0.083 | -0.652 | 0.514 | |
| | Cold-acclimated | *LacZ*- injected (intercept) | 2.513 | 0.084 | 29.842 | < 0.001 | 2,77 |
| | | No-injection | -0.275 | 0.103 | -2.667 | **0.008** | |
| | | *vrille*- injected | 0.261 | 0.106 | 2.454 | **0.014** | |

Table S12. The number of pairwise nucleotide differences in *vrille* exons among *D. montana* and *D. flavomontana* populations.

| | *D. fla* Ashford | *D. fla* Cranbrook | *D. fla* Jackson | *D. fla* Liberty | *D. fla* Livingston | *D. fla* McBride | *D. fla* Terrace | *D. fla* Vancouver | *D. mon* Afton | *D. mon* Ashford | *D. mon* Cranbrook | *D. mon* Jackson | *D. mon* McBride | *D. mon* Seward | *D. mon* Terrace | *D. mon* Vancouver |
|---|---|---|---|---|---|---|---|---|---|---|---|---|---|---|---|---|
| ***D. fla* Ashford** | 0 | 6 | 2 | 6 | 8 | 14 | 7 | 5 | 86 | 88 | 85 | 86 | 80 | 90 | 87 | 85 |
| ***D. fla* Cranbrook** | 6 | 0 | 6 | 8 | 6 | 10 | 11 | 7 | 82 | 84 | 81 | 82 | 76 | 86 | 83 | 81 |
| ***D. fla* Jackson** | 2 | 6 | 0 | 6 | 6 | 12 | 7 | 3 | 86 | 88 | 85 | 86 | 80 | 90 | 87 | 85 |
| ***D. fla* Liberty** | 6 | 8 | 6 | 0 | 12 | 16 | 13 | 7 | 84 | 86 | 83 | 84 | 78 | 88 | 85 | 83 |
| ***D. fla* Livingston** | 8 | 6 | 6 | 12 | 0 | 10 | 9 | 7 | 86 | 88 | 83 | 84 | 80 | 90 | 85 | 83 |
| ***D. fla* McBride** | 14 | 10 | 12 | 16 | 10 | 0 | 15 | 11 | 88 | 90 | 85 | 86 | 82 | 92 | 87 | 85 |
| ***D. fla* Terrace** | 7 | 11 | 7 | 13 | 9 | 15 | 0 | 8 | 89 | 93 | 88 | 91 | 85 | 93 | 90 | 88 |
| ***D. fla* Vancouver** | 5 | 7 | 3 | 7 | 7 | 11 | 8 | 0 | 83 | 85 | 82 | 83 | 77 | 87 | 84 | 82 |
| ***D. mon* Afton** | 86 | 82 | 86 | 84 | 86 | 88 | 89 | 83 | 0 | 18 | 20 | 12 | 22 | 25 | 21 | 22 |
| ***D. mon* Ashford** | 88 | 84 | 88 | 86 | 88 | 90 | 93 | 85 | 18 | 0 | 28 | 20 | 18 | 31 | 29 | 30 |
| ***D. mon* Cranbrook** | 85 | 81 | 85 | 83 | 83 | 85 | 88 | 82 | 20 | 28 | 0 | 12 | 20 | 27 | 23 | 14 |
| ***D. mon* Jackson** | 86 | 82 | 86 | 84 | 84 | 86 | 91 | 83 | 12 | 20 | 12 | 0 | 22 | 29 | 21 | 14 |
| ***D. mon* McBride** | 80 | 76 | 80 | 78 | 80 | 82 | 85 | 77 | 22 | 18 | 20 | 22 | 0 | 25 | 23 | 20 |
| ***D. mon* Seward** | 90 | 86 | 90 | 88 | 90 | 92 | 93 | 87 | 25 | 31 | 27 | 29 | 25 | 0 | 24 | 23 |
| ***D. mon* Terrace** | 87 | 83 | 87 | 85 | 85 | 87 | 90 | 84 | 21 | 29 | 23 | 21 | 23 | 24 | 0 | 21 |
| ***D. mon* Vancouver** | 85 | 81 | 85 | 83 | 83 | 85 | 88 | 82 | 22 | 30 | 14 | 14 | 20 | 23 | 21 | 0 |

Table S13. The number of pairwise amino acid differences in translated *vrille* exons among *D. montana* and *D. flavomontana* populations.

| | *D. fla* Ashford | *D. fla* Cranbrook | *D. fla* Jackson | *D. fla* Liberty | *D. fla* Livingston | *D. fla* McBride | *D. fla* Terrace | *D. fla* Vancouver | *D. mon* Afton | *D. mon* Ashford | *D. mon* Cranbrook | *D. mon* Jackson | *D. mon* McBride | *D. mon* Seward | *D. mon* Terrace | *D. mon* Vancouver |
|---|---|---|---|---|---|---|---|---|---|---|---|---|---|---|---|---|
| ***D. fla* Ashford** | 0 | 3 | 1 | 2 | 4 | 7 | 3 | 3 | 22 | 22 | 23 | 23 | 19 | 25 | 28 | 24 |
| ***D. fla* Cranbrook** | 3 | 0 | 2 | 3 | 1 | 4 | 4 | 2 | 19 | 19 | 20 | 20 | 16 | 22 | 25 | 21 |
| ***D. fla* Jackson** | 1 | 2 | 0 | 1 | 3 | 6 | 4 | 2 | 21 | 21 | 22 | 22 | 18 | 24 | 27 | 23 |
| ***D. fla* Liberty** | 2 | 3 | 1 | 0 | 4 | 7 | 5 | 3 | 22 | 22 | 23 | 23 | 19 | 25 | 28 | 24 |
| ***D. fla* Livingston** | 4 | 1 | 3 | 4 | 0 | 3 | 5 | 3 | 20 | 20 | 19 | 19 | 17 | 23 | 24 | 20 |
| ***D. fla* McBride** | 7 | 4 | 6 | 7 | 3 | 0 | 8 | 4 | 21 | 21 | 21 | 20 | 18 | 24 | 25 | 21 |
| ***D. fla* Terrace** | 3 | 4 | 4 | 5 | 5 | 8 | 0 | 4 | 21 | 23 | 22 | 24 | 20 | 25 | 27 | 23 |
| ***D. fla* Vancouver** | 3 | 2 | 2 | 3 | 3 | 4 | 4 | 0 | 19 | 19 | 21 | 20 | 16 | 22 | 25 | 21 |
| ***D. mon* Afton** | 22 | 19 | 21 | 22 | 20 | 21 | 21 | 19 | 0 | 6 | 6 | 5 | 5 | 10 | 10 | 6 |
| ***D. mon* Ashford** | 22 | 19 | 21 | 22 | 20 | 21 | 23 | 19 | 6 | 0 | 10 | 7 | 3 | 9 | 14 | 10 |
| ***D. mon* Cranbrook** | 23 | 20 | 22 | 23 | 19 | 21 | 22 | 21 | 6 | 10 | 0 | 3 | 7 | 12 | 8 | 4 |
| ***D. mon* Jackson** | 23 | 20 | 22 | 23 | 19 | 20 | 24 | 20 | 5 | 7 | 3 | 0 | 6 | 12 | 9 | 5 |
| ***D. mon* McBride** | 19 | 16 | 18 | 19 | 17 | 18 | 20 | 16 | 5 | 3 | 7 | 6 | 0 | 6 | 11 | 7 |
| ***D. mon* Seward** | 25 | 22 | 24 | 25 | 23 | 24 | 25 | 22 | 10 | 9 | 12 | 12 | 6 | 0 | 14 | 12 |
| ***D. mon* Terrace** | 28 | 25 | 27 | 28 | 24 | 25 | 27 | 25 | 10 | 14 | 8 | 9 | 11 | 14 | 0 | 10 |
| ***D. mon* Vancouver** | 24 | 21 | 23 | 24 | 20 | 21 | 23 | 21 | 6 | 10 | 4 | 5 | 7 | 12 | 10 | 0 |

Table S14. Pairwise differences in the mean $CT_{min}$ among *D. montana* and *D. flavomontana* populations.

| | ***D. fla*** **Ashford** | ***D. fla*** **Cranbrook** | ***D. fla*** **Jackson** | ***D. fla*** **Liberty** | ***D. fla*** **Livingston** | ***D. fla*** **McBride** | ***D. fla*** **Terrace** | ***D. fla*** **Vancouver** | ***D. mon*** **Afton** | ***D. mon*** **Ashford** | ***D. mon*** **Cranbrook** | ***D. mon*** **Jackson** | ***D. mon*** **McBride** | ***D. mon*** **Seward** | ***D. mon*** **Terrace** | ***D. mon*** **Vancouver** |
|---|---|---|---|---|---|---|---|---|---|---|---|---|---|---|---|---|
| ***D. fla*** **Ashford** | 0.00 | 0.19 | 0.03 | 0.12 | 0.33 | 0.10 | 0.25 | 0.24 | 1.55 | 1.65 | 1.60 | 1.68 | 1.66 | 1.80 | 1.34 | 1.41 |
| ***D. fla*** **Cranbrook** | 0.19 | 0.00 | 0.15 | 0.06 | 0.14 | 0.09 | 0.07 | 0.06 | 1.36 | 1.46 | 1.42 | 1.49 | 1.47 | 1.62 | 1.15 | 1.22 |
| ***D. fla*** **Jackson** | 0.03 | 0.15 | 0.00 | 0.09 | 0.30 | 0.06 | 0.22 | 0.21 | 1.51 | 1.62 | 1.57 | 1.64 | 1.63 | 1.77 | 1.30 | 1.38 |
| ***D. fla*** **Liberty** | 0.12 | 0.06 | 0.09 | 0.00 | 0.21 | 0.03 | 0.13 | 0.12 | 1.42 | 1.53 | 1.48 | 1.55 | 1.54 | 1.68 | 1.21 | 1.28 |
| ***D. fla*** **Livingston** | 0.33 | 0.14 | 0.30 | 0.21 | 0.00 | 0.23 | 0.07 | 0.08 | 1.22 | 1.32 | 1.27 | 1.35 | 1.33 | 1.47 | 1.01 | 1.08 |
| ***D. fla*** **McBride** | 0.10 | 0.09 | 0.06 | 0.03 | 0.23 | 0.00 | 0.16 | 0.15 | 1.45 | 1.55 | 1.51 | 1.58 | 1.56 | 1.70 | 1.24 | 1.31 |
| ***D. fla*** **Terrace** | 0.25 | 0.07 | 0.22 | 0.13 | 0.07 | 0.16 | 0.00 | 0.01 | 1.29 | 1.40 | 1.35 | 1.42 | 1.40 | 1.55 | 1.08 | 1.15 |
| ***D. fla*** **Vancouver** | 0.24 | 0.06 | 0.21 | 0.12 | 0.08 | 0.15 | 0.01 | 0.00 | 1.30 | 1.41 | 1.36 | 1.43 | 1.41 | 1.56 | 1.09 | 1.16 |
| ***D. mon*** **Afton** | 1.55 | 1.36 | 1.51 | 1.42 | 1.22 | 1.45 | 1.29 | 1.30 | 0.00 | 0.10 | 0.06 | 0.13 | 0.11 | 0.26 | 0.21 | 0.14 |
| ***D. mon*** **Ashford** | 1.65 | 1.46 | 1.62 | 1.53 | 1.32 | 1.55 | 1.40 | 1.41 | 0.10 | 0.00 | 0.05 | 0.03 | 0.01 | 0.15 | 0.31 | 0.24 |
| ***D. mon*** **Cranbrook** | 1.60 | 1.42 | 1.57 | 1.48 | 1.27 | 1.51 | 1.35 | 1.36 | 0.06 | 0.05 | 0.00 | 0.07 | 0.06 | 0.20 | 0.27 | 0.20 |
| ***D. mon*** **Jackson** | 1.68 | 1.49 | 1.64 | 1.55 | 1.35 | 1.58 | 1.42 | 1.43 | 0.13 | 0.03 | 0.07 | 0.00 | 0.02 | 0.13 | 0.34 | 0.27 |
| ***D. mon*** **McBride** | 1.66 | 1.47 | 1.63 | 1.54 | 1.33 | 1.56 | 1.40 | 1.41 | 0.11 | 0.01 | 0.06 | 0.02 | 0.00 | 0.14 | 0.32 | 0.25 |
| ***D. mon*** **Seward** | 1.80 | 1.62 | 1.77 | 1.68 | 1.47 | 1.70 | 1.55 | 1.56 | 0.26 | 0.15 | 0.20 | 0.13 | 0.14 | 0.00 | 0.46 | 0.39 |
| ***D. mon*** **Terrace** | 1.34 | 1.15 | 1.30 | 1.21 | 1.01 | 1.24 | 1.08 | 1.09 | 0.21 | 0.31 | 0.27 | 0.34 | 0.32 | 0.46 | 0.00 | 0.07 |
| ***D. mon*** **Vancouver** | 1.41 | 1.22 | 1.38 | 1.28 | 1.08 | 1.31 | 1.15 | 1.16 | 0.14 | 0.24 | 0.20 | 0.27 | 0.25 | 0.39 | 0.07 | 0.00 |

Table S15. Pairwise differences in the mean CCRT among *D. montana* and *D. flavomontana* populations.

| | ***D. fla*** **Ashford** | ***D. fla*** **Cranbrook** | ***D. fla*** **Jackson** | ***D. fla*** **Liberty** | ***D. fla*** **Livingston** | ***D. fla*** **McBride** | ***D. fla*** **Terrace** | ***D. fla*** **Vancouver** | ***D. mon*** **Afton** | ***D. mon*** **Ashford** | ***D. mon*** **Cranbrook** | ***D. mon*** **Jackson** | ***D. mon*** **McBride** | ***D. mon*** **Seward** | ***D. mon*** **Terrace** | ***D. mon*** **Vancouver** |
|---|---|---|---|---|---|---|---|---|---|---|---|---|---|---|---|---|
| ***D. fla*** **Ashford** | 0.00 | 1.03 | 0.93 | 0.88 | 1.01 | 1.67 | 4.36 | 2.56 | 4.81 | 7.18 | 6.89 | 7.03 | 7.15 | 7.73 | 7.37 | 7.35 |
| ***D. fla*** **Cranbrook** | 1.03 | 0.00 | 0.10 | 0.15 | 0.02 | 0.63 | 3.33 | 1.52 | 3.78 | 6.15 | 5.86 | 6.00 | 6.12 | 6.70 | 6.34 | 6.32 |
| ***D. fla*** **Jackson** | 0.93 | 0.10 | 0.00 | 0.04 | 0.08 | 0.74 | 3.43 | 1.63 | 3.88 | 6.25 | 5.96 | 6.10 | 6.22 | 6.80 | 6.44 | 6.43 |
| ***D. fla*** **Liberty** | 0.88 | 0.15 | 0.04 | 0.00 | 0.13 | 0.78 | 3.47 | 1.67 | 3.92 | 6.29 | 6.01 | 6.15 | 6.27 | 6.85 | 6.49 | 6.47 |
| ***D. fla*** **Livingston** | 1.01 | 0.02 | 0.08 | 0.13 | 0.00 | 0.65 | 3.35 | 1.54 | 3.80 | 6.17 | 5.88 | 6.02 | 6.14 | 6.72 | 6.36 | 6.34 |
| ***D. fla*** **McBride** | 1.67 | 0.63 | 0.74 | 0.78 | 0.65 | 0.00 | 2.69 | 0.89 | 3.14 | 5.51 | 5.23 | 5.36 | 5.49 | 6.07 | 5.70 | 5.69 |
| ***D. fla*** **Terrace** | 4.36 | 3.33 | 3.43 | 3.47 | 3.35 | 2.69 | 0.00 | 1.80 | 0.45 | 2.82 | 2.53 | 2.67 | 2.79 | 3.37 | 3.01 | 2.99 |
| ***D. fla*** **Vancouver** | 2.56 | 1.52 | 1.63 | 1.67 | 1.54 | 0.89 | 1.80 | 0.00 | 2.25 | 4.62 | 4.34 | 4.48 | 4.60 | 5.18 | 4.81 | 4.80 |
| ***D. mon*** **Afton** | 4.81 | 3.78 | 3.88 | 3.92 | 3.80 | 3.14 | 0.45 | 2.25 | 0.00 | 2.37 | 2.08 | 2.22 | 2.34 | 2.92 | 2.56 | 2.54 |
| ***D. mon*** **Ashford** | 7.18 | 6.15 | 6.25 | 6.29 | 6.17 | 5.51 | 2.82 | 4.62 | 2.37 | 0.00 | 0.29 | 0.15 | 0.03 | 0.55 | 0.19 | 0.17 |
| ***D. mon*** **Cranbrook** | 6.89 | 5.86 | 5.96 | 6.01 | 5.88 | 5.23 | 2.53 | 4.34 | 2.08 | 0.29 | 0.00 | 0.14 | 0.26 | 0.84 | 0.48 | 0.46 |
| ***D. mon*** **Jackson** | 7.03 | 6.00 | 6.10 | 6.15 | 6.02 | 5.36 | 2.67 | 4.48 | 2.22 | 0.15 | 0.14 | 0.00 | 0.12 | 0.70 | 0.34 | 0.32 |
| ***D. mon*** **McBride** | 7.15 | 6.12 | 6.22 | 6.27 | 6.14 | 5.49 | 2.79 | 4.60 | 2.34 | 0.03 | 0.26 | 0.12 | 0.00 | 0.58 | 0.22 | 0.20 |
| ***D. mon*** **Seward** | 7.73 | 6.70 | 6.80 | 6.85 | 6.72 | 6.07 | 3.37 | 5.18 | 2.92 | 0.55 | 0.84 | 0.70 | 0.58 | 0.00 | 0.36 | 0.38 |
| ***D. mon*** **Terrace** | 7.37 | 6.34 | 6.44 | 6.49 | 6.36 | 5.70 | 3.01 | 4.81 | 2.56 | 0.19 | 0.48 | 0.34 | 0.22 | 0.36 | 0.00 | 0.02 |
| ***D. mon*** **Vancouver** | 7.35 | 6.32 | 6.43 | 6.47 | 6.34 | 5.69 | 2.99 | 4.80 | 2.54 | 0.17 | 0.46 | 0.32 | 0.20 | 0.38 | 0.02 | 0.00 |

Table S16. Quantitative real time PCR (qPCR) primers and their efficiencies (%) for *vrille* gene and reference genes (*Tub2*, *Rpl32*), and primers designed for dsRNA used in RNA interference (RNAi). Primers were designed based on *D. montana* genomic sequences under NCBI accession number LUVX00000000 [3] together with information from *D. virilis* exons (Flybase) using Primer3 (primer3.ut.ee) and NetPrimer (www.premierbiosoft.com/ netprimer) programs.

| Primers | Gene | Source | Forward primer sequence | Reverse primer sequence | Efficiency value (E%) for qPCR primers |
|---|---|---|---|---|---|
| **qPCR** | *vrille* | Flybase (FBgn0204709) | 5’ – CTTTTTTCAAGAGACTGGAACTACT – 3’ | 5’ – GCACTGCGTATGTAGAATGTTG – 3’ | 98.1 |
| | *Tubulin beta chain* (*Tub2*) | Flybase (FBgn0208711) | 5’ – CGTGCTGTGTTTGTCGATCT– 3’ | 5’ –GATCTCCTTGCCAATGGTGT– 3’ | 96.0 |
| | *Ribosomal Protein L32* (*RpL32*) | Flybase (FBgn0016459) | 5’ – CATCAGCAGCACCTCCAGTTC – 3’ | 5’ – GATATGCCAAGCTGTCGCACAA – 3’ | 97.8 |
| **dsRNA synthesis** | *vrille* | Flybase (FBgn0204709) | 5’ GCGATATACATATGGTGAATGAAGTG 3’ | 5’ GCTCCAACGGGCTTTCTATC 3’ | - |
| | *LacZ* | [4] | 5’ AGAATCCGACGGGTTGTTACT 3’ | 5’ CACCACGCTCATCGATAATTT 3’ | - |
| | pJET 1.2 F / pJET 1.2. T7 R | CloneJET PCR Cloning Kit (Thermo Fisher Scientific) | 5’ –CGACTCACTATAGGGAGAGCGGC – 3’ | 5’ – AAGAACATCGATTTTCCATGGCAG – 3’ | - |

### Supplementary figures

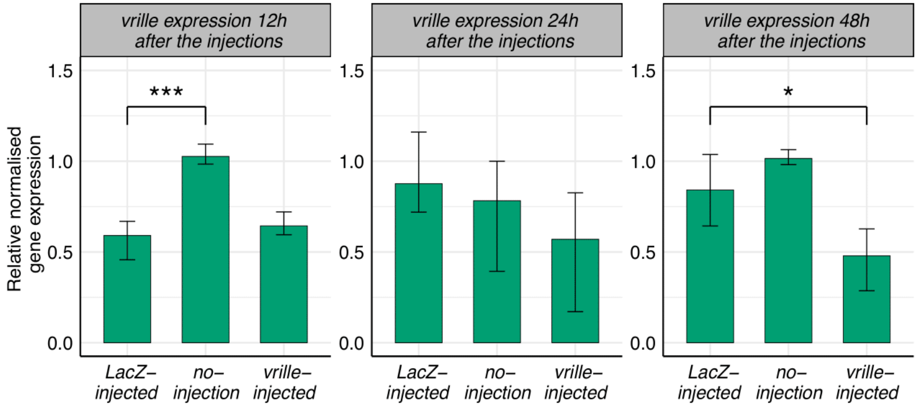


| Response time | Treatment | Coefficient | SE | t | P |
|---|---|---|---|---|---|
| 12h | *LacZ*- injected (intercept) | 0.742 | 0.062 | 12.045 | < 0.001 |
| | No-injection | 0.547 | 0.087 | 6.277 | **< 0.001** |
| | *vrille* -injected | 0.067 | 0.087 | 0.766 | 0.473 |
| 24h | *LacZ*- injected (intercept) | 4.031 | 0.837 | 4.817 | 0.003 |
| | No-injection | -0.431 | 1.184 | -0.364 | 0.728 |
| | *vrille*- injected | -1.409 | 1.184 | -1.191 | 0.279 |
| 48h | *LacZ*- injected (intercept) | 1.208 | 0.128 | 9.467 | < 0.001 |
| | No-injection | 0.249 | 0.180 | 1.378 | 0.218 |
| | *vrille*- injected | -0.521 | 0.180 | -2.888 | **0.028** |

Figure S1. The effectiveness of RNAi was investigated 12, 24 and 48 hours after injections. Expression levels of *vrille* were compared between *LacZ*-injected females and no-injection and *vrille*-injected females. Error bars represent bootstrapped 95% confidence intervals. Significance levels were obtained from a linear model (ANOVA) and only significant differences are shown: * P < 0.05, ** P < 0.01 and *** P < 0.001.

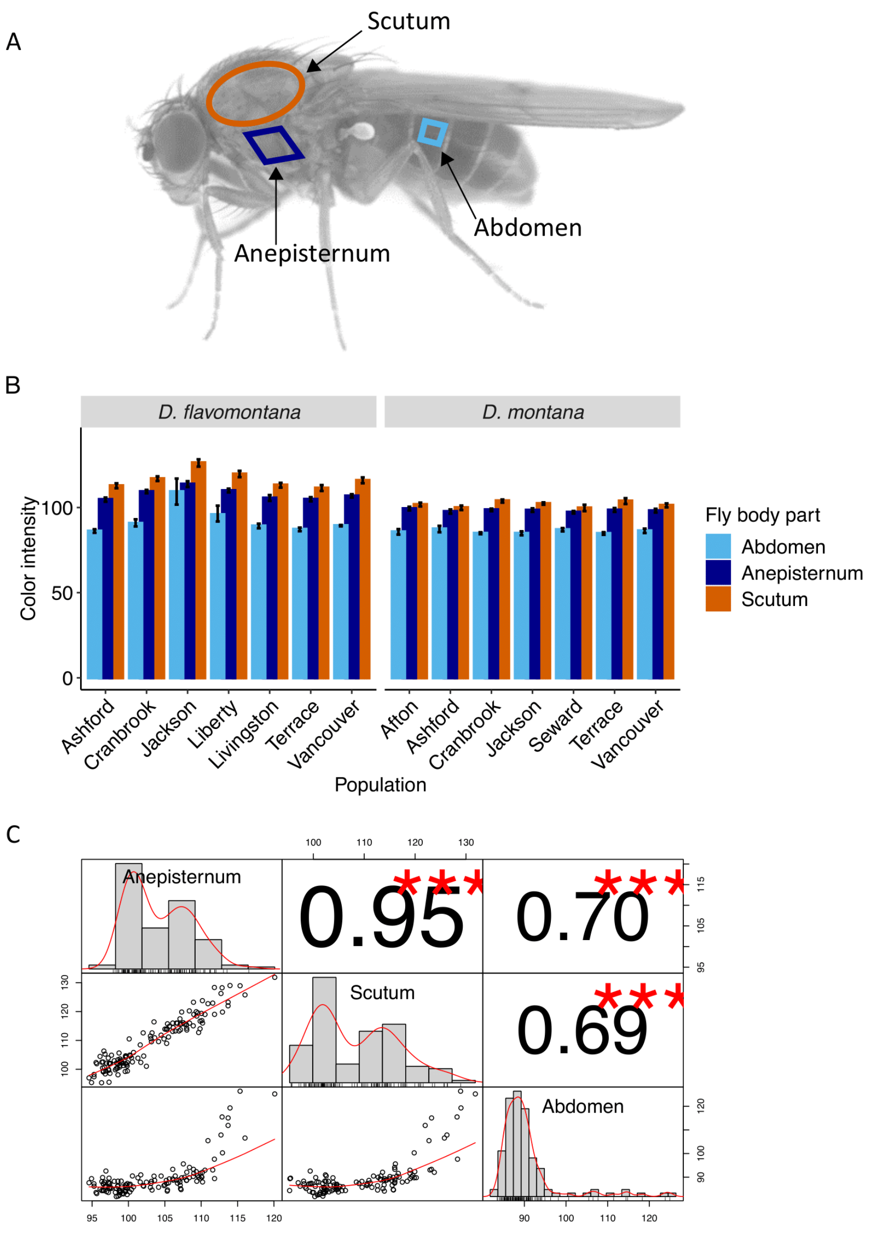


Figure S2. Preliminary colour intensity measurements of different *D. montana* and *D. flavomontana* populations. (A) Colour measurements were taken from two parts of the thorax, scutum and anepisternum, and from A3 segment of the abdomen. The photo of *D. flavomontana* was taken by Noora Poikela. (B) Measurements were taken from 5 females per strain, and 1-2 strains per population in each species (McBride populations were not included), and linearly scaled from 0 to 255 (0 = black, 255 = white). Scutum and anepisternum incorporated most of the colour intensity variation among *D. montana* and *D. flavomontana* flies, while abdomen was equally dark among them, except in *D. flavomontana* from Jackson and Liberty populations, which showed slightly more variation. Error bars represent bootstrapped 95% confidence intervals. (C) Scutum and anepisternum were highly correlated with each other (Pearson correlation coefficient = 0.95), enabling us to use only the former in our colour analysis.

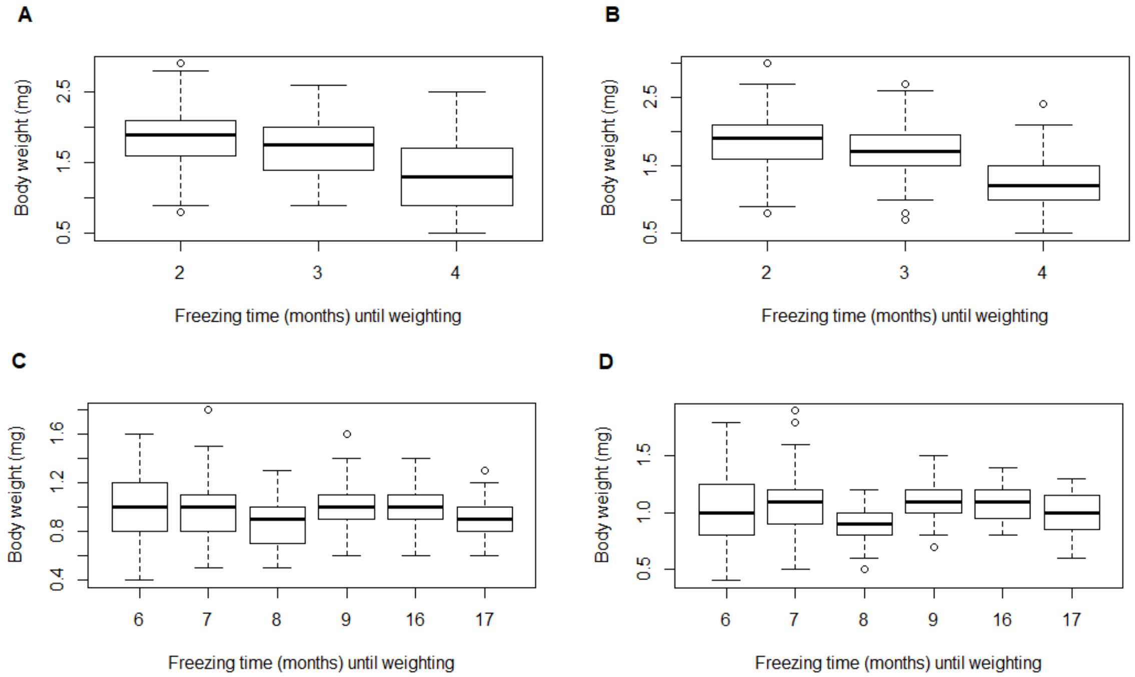


Fig. S3. The effect of the freezing time (in months) on body weight in (A) $CT_{min}$ *D. montana* flies (GLMM, $z_{2,443}$=-5.556, P<0.001), (B) $CT_{min}$ *D. flavomontana* flies (GLMM, $z_{2,334}$= -6.437, P<0.001), (C) CCRT *D. montana* flies (GLMM, $z_{2,437}$=-0.900, P=0.368) and (D) CCRT *D. flavomontana* flies (GLMM, $z_{2,437}$=-0.645, P=0.519).

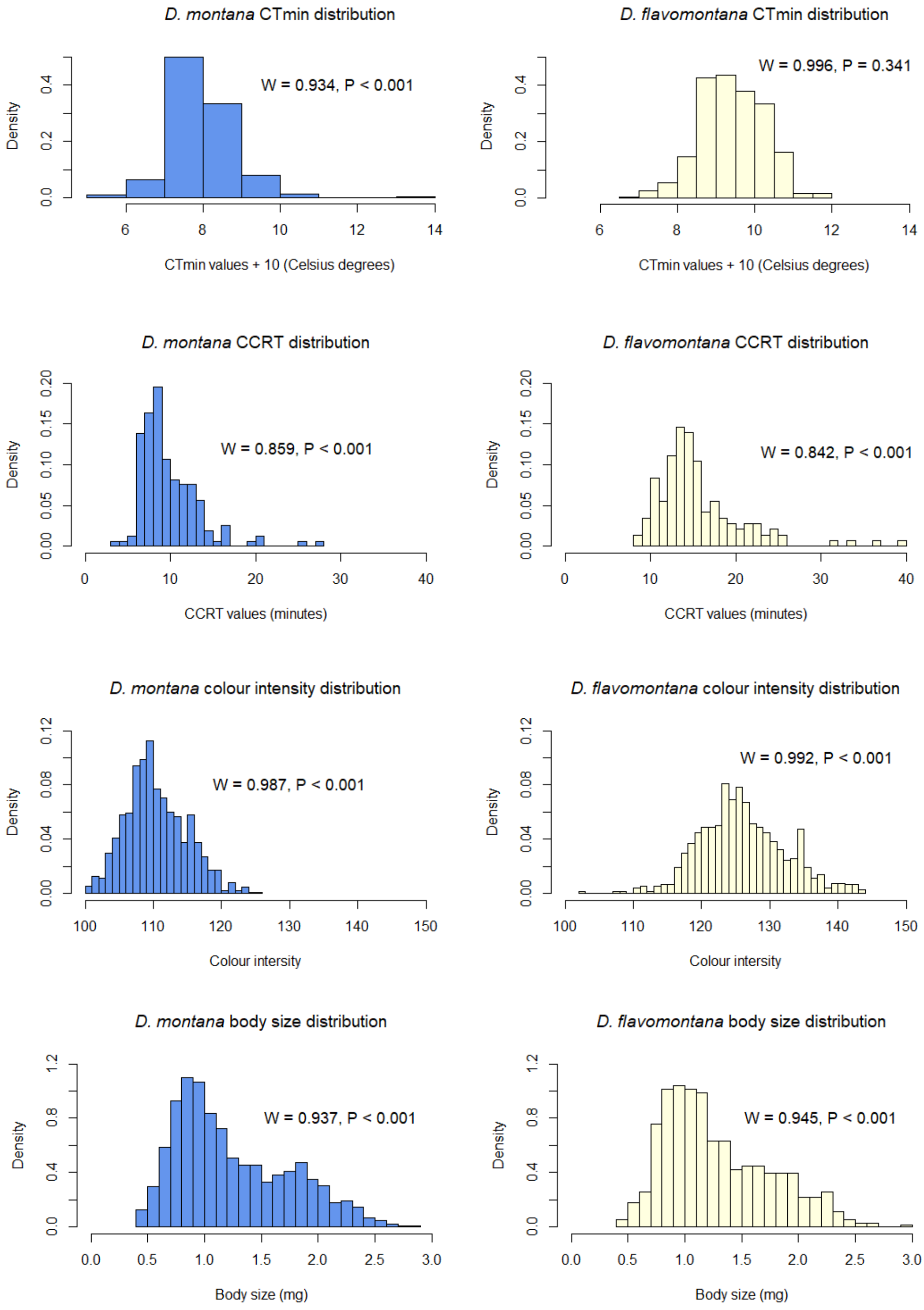


Figure S4. Distributions and Shapiro-Wilk test statistics and P-values for testing normality of $CT_{min}$, CCRT, body colour and body size (measured as weight) data of *D. montana* and *D. flavomontana*.

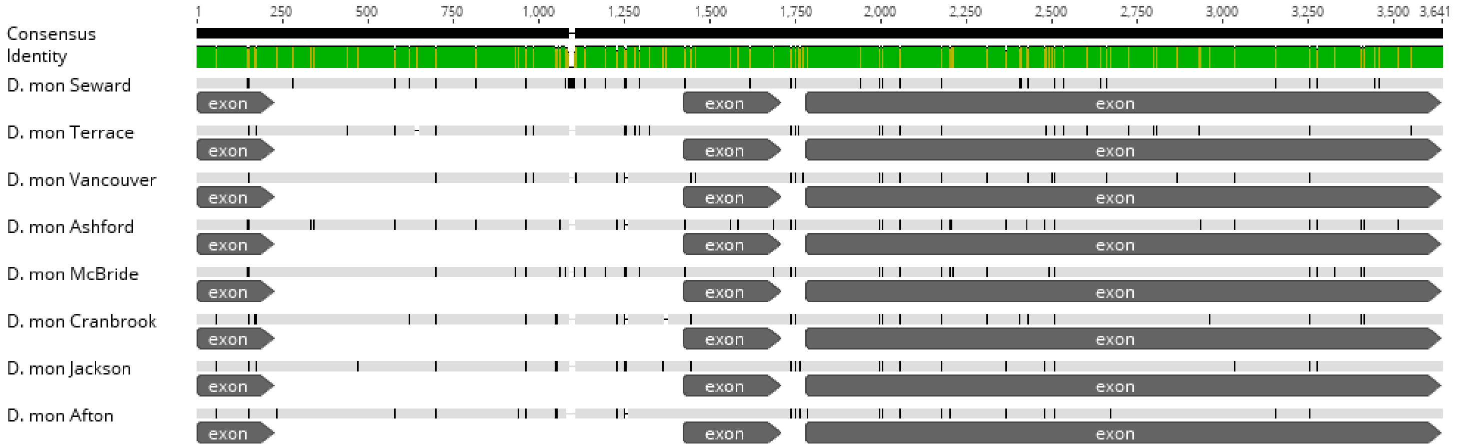


Figure S5. *vrille* sequence alignment and annotation of eight *D. montana* samples originating across study sites.

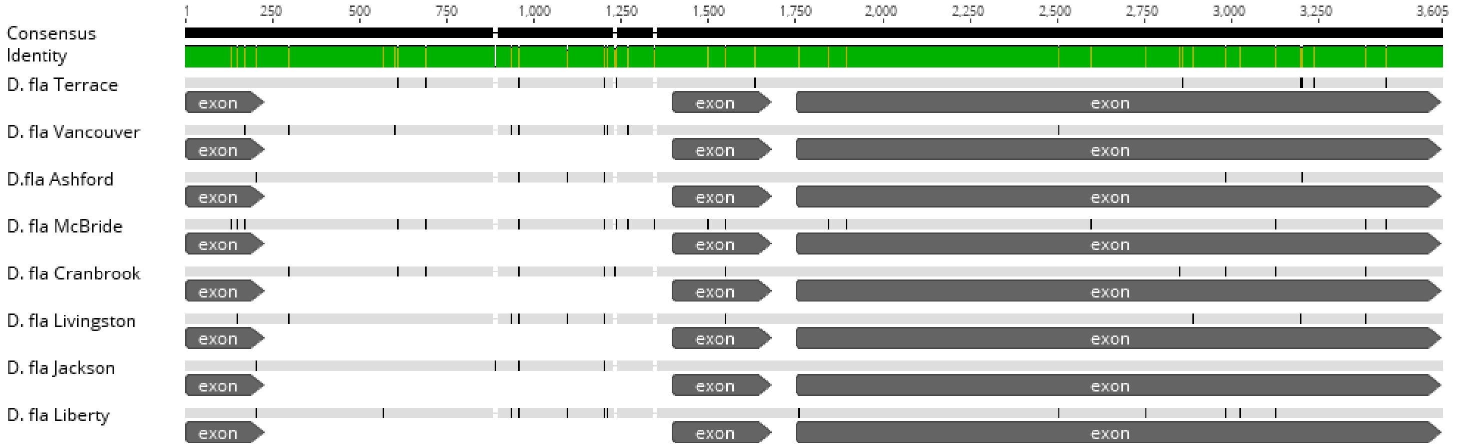


Figure S6. *vrille* sequence alignment and annotation of eight *D. flavomontana* samples originating across study sites.

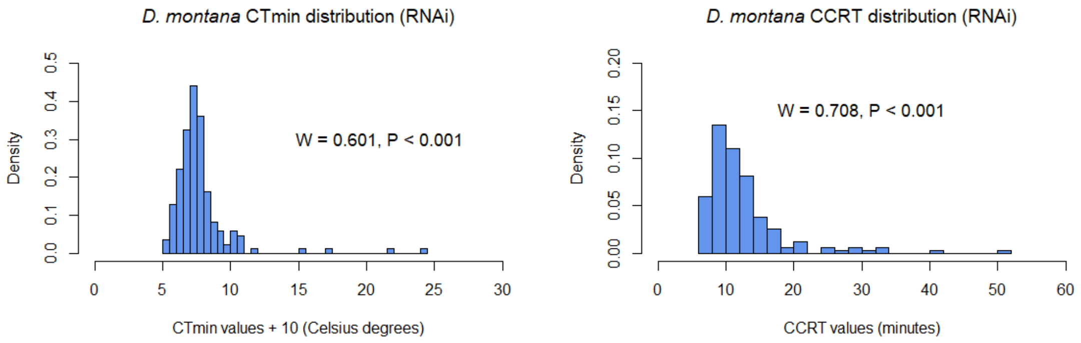


Figure S7. Distributions and Shapiro-Wilk test statistics and P-values for testing normality of $CT_{min}$ and CCRT data of *D. montana* in RNAi studies.

#### Supplementary references

1. Fick SE, Hijmans RJ. WorldClim 2: new 1-km spatial resolution climate surfaces for global land areas. Int J Climatol. 2017;37:4302–15.

2. Tyukmaeva VI, Lankinen P, Kinnunen J, Kauranen H, Hoikkala A. Latitudinal clines in the timing and temperature-sensitivity of photoperiodic reproductive diapause in *Drosophila montana*. Ecography (Cop). 2020;43:1–10.

3. Parker DJ, Wiberg RAW, Trivedi U, Tyukmaeva VI, Gharbi K, Butlin RK, et al. Inter and intraspecific genomic divergence in *Drosophila montana* shows evidence for cold adaptation. Genome Biol Evol. 2018;10:2086–101.

4. Vigoder, F. M., Parker, D. J., Cook, N., Tournière, O., Sneddon, T., & Ritchie, M. G. (2016). Inducing Cold-Sensitivity in the Frigophilic Fly *Drosophila montana* by RNAi. PLoS ONE, 11(11), 1–9. doi: 10.1371/journal.pone.0165724
